# Modular Assembly of an Infectious Clone of the Akata strain of EBV using Synthetic Genomics Methods in Yeast

**DOI:** 10.64898/2026.06.16.732627

**Authors:** Adam Raviv, Kristiana Smith, Shuang Chang, Sanjana Prasad, Peter Grzesik, Sydney Ghoreishi, Bogdan Paun, Lauren Oldfield, Angelica Contreras, Jennifer Petr, Dirk Dittmer, Rena Xian, Gabriel Ghiaur, Sanjay Vashee, Richard F. Ambinder, Prashant J. Desai

## Abstract

We first showed in 2015 the use of synthetic biology recombination methods in yeast to build herpes simplex virus type-1 (HSV-1) and human cytomegalovirus (HCMV) genomes from multiple fragments. The genomes were built using transformation-associated recombination (TAR) in yeast, by virtue of overlapping sequences between the different fragments. In the present study we demonstrate the successful assembly of the Epstein-Barr virus (EBV) genome using similar methods. We used as the model genome, the Akata Burkitt’s lymphoma genome, specifically the BX1 genome which encodes a neomycin selectable marker and a GFP expression cassette in the *BXLF1* region. The 171.3 kb genome was first deconstructed into 11 fragments *in silico*, each having 79 bp overlapping sequence between the fragments. The 11 fragments (TAR 1 to TAR 11) were cloned using TAR in yeast, analyzed by restriction enzyme analyses and Nanopore sequencing to validate the cloned fragment. The EBV genome was built in two stages: TAR fragments 1 to 6 and TAR fragments 7 to 11 were assembled to generate two half-genomes. The whole genome (TAR 1-11) was then assembled by joining TAR 1-6 with TAR 7-11. EBV yeast-assembled genomes were examined by PCR assays, restriction enzyme analyses and sequencing and transfected into HEK-293 cells to generate virus producer cell lines. The HEK-293 cell clones were tested for virus production following lytic induction using a novel baculovirus transduction method for expression of Zta, Rta and glycoprotein B (*BALF4*). The supernatants from these induced cells were harvested and used to infect Raji cells. This analysis revealed a significant number of cells displaying strong GFP fluorescence indicative of infectious virus. We used this supernatant virus to infect primary B cells and were able to derive lymphoblastoid cell lines (LCL) indicative of the ability of this virus to transform B cells. We tested this method for engineering different mutations. Two mutations were made, one in *Zta* and the other in the small capsid protein (*BFRF3*). Mutations were engineered in the TAR plasmid in which the genes reside and after sequence validation, assembled into the TAR 1-6 half genome and then the TAR 1-11 genome, which was used to generate HEK-293 cell clones. For the ΔZta cell lines, we could detect virus in the supernatants only if baculovirus expressing Zta *in trans* was included, this ΔZta EBV could transform B cells. The small capsid protein (BFRF3) decorates the capsid shell and is required for EBV capsid assembly in a self-assembly system. When the HEK-293 cell clones were induced using co-expression of Zta, Rta and gB, no virus was detected in the culture supernatants. However, if we provided BFRF3 *in trans* using transduction with a baculovirus expressing this protein, virus was detected in the supernatants. This provides the first report of the essential role of the small capsid protein in EBV-infected cells.

## INTRODUCTION

Herpesviruses are large double-stranded DNA viruses that are disseminated throughout the animal kingdom. They are major human pathogens, they cause life-long persistent infections with clinical manifestations ranging from mild cold sores to lymphomas and angiogenic tumors. The large genomes of herpesviruses have a coding capacity exceeding 100 genes in some cases. The ability to manipulate these genomes and subsequently assign functions to the gene products within these large elements was advanced by bacterial artificial chromosome (BAC) technology in *Escherichia coli (E.coli)* [1-29]. We have developed an alternative approach to herpesvirus genome engineering by using novel synthetic genomics methods in yeast. We previously demonstrated the ability to build infectious clones of herpes simplex virus type 1 (HSV-1) [30] and human cytomegalovirus (HCMV) genomes [31]. These studies were used to establish and refine synthetic genomics engineering methods; we then applied this information to build an infectious clone of the Epstein-Barr virus (EBV) that is stable and can be engineered using synthetic biology.

EBV, a member of the gammaherpesvirus family, primarily infects B lymphocytes and epithelial cells, and is responsible for several lymphoproliferative diseases [32-36]. Primary EBV infection in adolescents often results in infectious mononucleosis (IM). EBV is also associated with malignancies, such as Burkitt’s lymphoma (BL), nasopharyngeal carcinoma and post-transplant lymphoproliferative disease, and it has more recently been implicated in multiple sclerosis [37-39]. The EBV Akata genome is 171.3 kb in length, containing multiple complex functional repeat regions (Fig. 1) [40]. There are variable terminal repeat (TR) regions in the circular genome [41], four internal repeat (IR) regions labeled IR1 to IR4, as well as three origins of replication (oriP, oriLyt-L and R) [42]. IR1 contains the largest repeat array composed of multiple copies of the BamHI-W (BamW) sequence. It encodes the leader protein exon of Epstein-Barr nuclear antigen (EBNA) and the Wp latency promoter. IR2 and IR4 are G-rich repetitive sequences that are located next to the lytic origins of replication, oriLyt L and R sequences, respectively. IR3 comprises the repetitive sequence encoding amino acids Gly-Ala within the EBNA1 protein. There is also the important family of repeats (FR) in the oriP region which is important for maintenance of the episome in infected cells. Finally, there are also repeat sequences within the coding regions of EBNA2, EBNA3B and EBNA3C that encode repetitive amino acids sequences [34, 43]. The EBV B95-8 strain is highly inducible for lytic replication [44] and was the first EBV genome sequenced and cloned as a BAC plasmid [13, 45, 46]. However, it has a 12 kb deletion, that includes the LF open reading frames, the duplicated DS-R copy of oriLyt and the BART (BamHI A rightward transcripts) miRNAs locus [47, 48]. The B95-8 BAC has been used extensively and successfully for selected knock-out studies, see [9] and references therein. A BAC clone with this 12 kb deletion repaired has also been generated [49]. We selected the Akata strain as the prototype EBV genome for synthetic genomics cloning. Because it has been sequenced [40] and a GFP-tagged recombinant, Akata-BX1 [50-52] can be induced using a more physiological IgG receptor antibody cross-linking approach, it is a suitable candidate for cloning by synthetic genomics [53]. Akata-BX1 refers to the virus originally modified by the Hutt-Fletcher Lab [52], which contains a CMV:GFP—HSV TK^Pr^:Neomycin cassette insertion within the non-essential *BXLF1* (EBV thymidine kinase) locus. This cassette allows for selection and visualization of cell lines harboring the genome [52]. This is a property of this virus that has proved useful. The normal Akata cell line loses the EBV genome after multiple passages [54]. Akata-BX1 cells retain the genome at higher copy number because of the selection pressure from replication in the presence of G418. While the Akata genome has been cloned using BAC technology in *E.coli*, it has in some instances shown instability in the family of repeats (FR) region [8, 55]. These repeats are essential for episomal maintenance of EBV in latently infected cells [56, 57]. CRISPR/Cas9 technology has also been used to clone out genomes from different EBV+ cell lines; notably two cloned genomes from SNU-719 and YCCEL1 remained intact as confirmed by long read sequencing [58]. Additionally, the M81 strain of EBV has been cloned as a BAC plasmid [10]. Recently, the Johannsen lab developed a dual-fluorescent reporter M81 recombinant (DFLR) that expresses green lantern fluorescent protein from the thymidine kinase (*BXLF1*) promoter and mScarlet fluorescent protein from the late *BILF2* promoter [59].

**Fig. 1.**
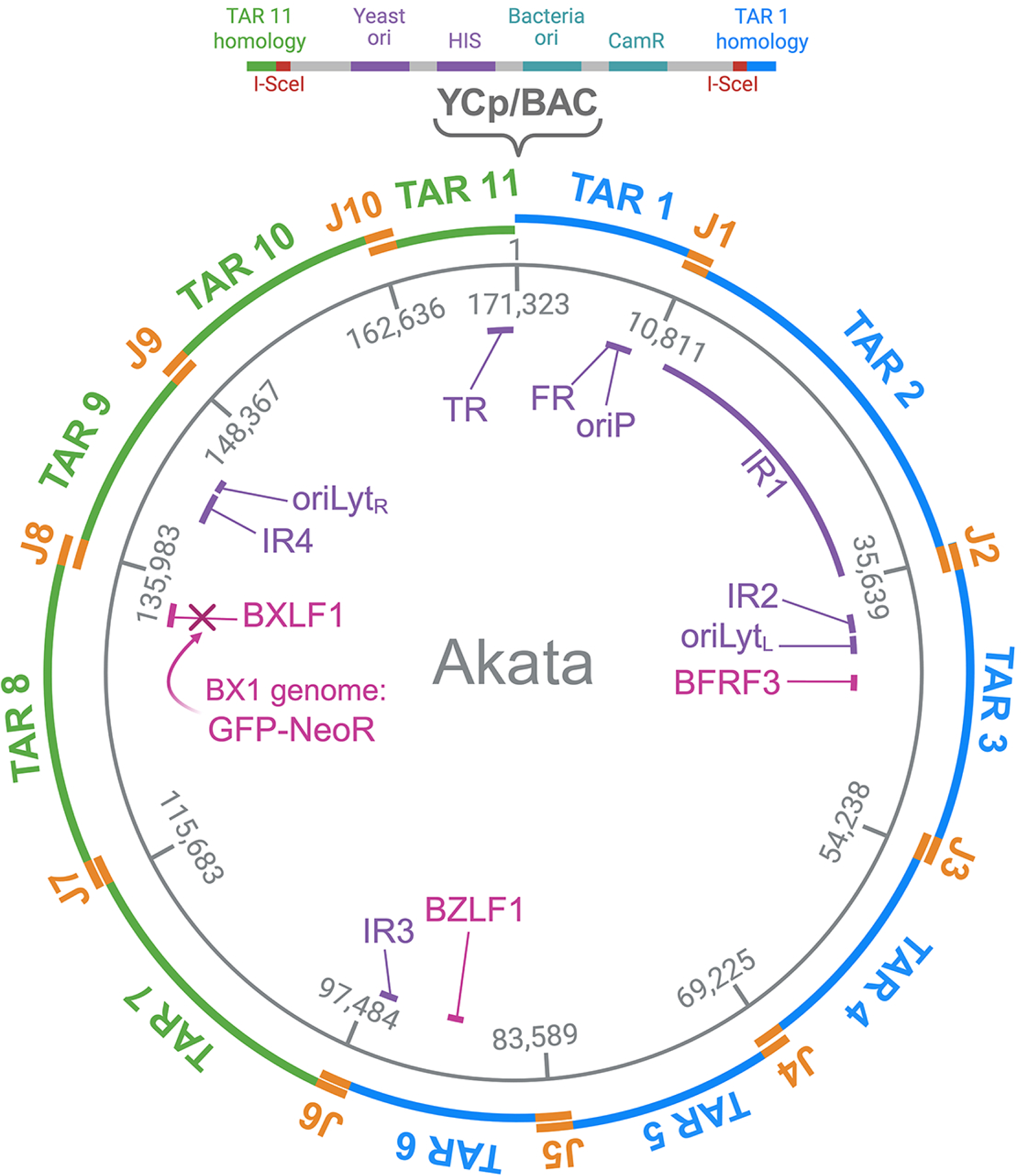
Design for the modular assembly of the EBV Akata-BX1 genome by transformation-associated recombination (TAR) cloning. The 171.3 kb EBV Akata genome [40] was divided *in silico* into 11 overlapping TAR fragments. The overlapping sequences between adjacent fragments are designated as junctions (J1-J10 shown in orange). Genome coordinates for each fragment are indicated. The BX1 genome encodes a GFP-Neomycin cassette inserted into the thymidine kinase (*BXLF1*) gene [52]. The blue and green TAR fragments were initially assembled into half-genomes (TAR 1-6 and TAR 7-11, respectively) before final assembly into the complete genome (TAR 1-11), where the YCp/BAC vector is positioned at the genomic ends between TAR 11 (after the TR region) and TAR 1. The YCp/BAC vector encodes TAR 11 homology (green) and TAR 1 homology (blue). The vector also encodes yeast and bacteria origins (ori) of replication and the histidine (HIS) nutritional selection marker and chloramphenicol resistance gene (CAM). Color coding: genes of interest (fuscia), repeat sequences (purple) and genome coordinates (grey).

In this study, we demonstrate that we can assemble the Akata genome carrying the BX1 cassette from 11 separate overlapping fragments in yeast using transformation-associated recombination (TAR) [60-62]. We show that these genomes can be propagated in HEK-293 cells and induced to produce virus using a novel baculovirus transduction method. Virus produced from these cells infects primary B cells, which eventually transform into lymphoblastoid cell lines (LCLs). Furthermore, we engineered null mutations in both the lytic transactivator, *Zta*, and the small capsid protein encoded by the *BFRF3* gene. These mutants display lethal phenotypes but can be rescued by expressing the respective gene *in trans* using baculovirus transduction. This work also provides the first demonstration that the EBV small capsid protein is essential for EBV virion formation.

## METHODS

### Cells and Viruses

Akata, Akata-BX1, Raji, primary B cells and lymphoblastoid cell lines (LCLs) cells were grown in RPMI medium supplemented with 10% fetal bovine serum (FBS; Invitrogen) and penicillin-streptomycin-neomycin (PSN) antibiotic mixture (Invitrogen). Cell lines were routinely passaged every 2-3 days using a 1:5 split ratio. HEK-293 and HEK-293-EBV^YA (yeast assembled)^ cells lines were cultured in high glucose DMEM containing 10% FBS and PSN and passaged every 2-3 days (2x10^6^ - 4x10^6^ cells per T75 flask). Both Akata-BX1 and HEK-293-EBV^YA^ cells were maintained in culture media containing 0.5 mg/ml Geneticin (Invitrogen). Insect Sf21 cells were passaged in Grace’s insect cell medium (Invitrogen) supplemented with 10% FBS and PSN and were passaged every 2 days (1x10^6^ cells/ml).

### Primary B cells

Primary B cells were isolated from bone marrow samples that were obtained as excess material from the harvests of normal donors for allogeneic bone marrow transplantation. Specimens were collected by the Johns Hopkins Kimmel Cancer Center Specimen Accessioning Core. See Ethical approval statement.

### Strains

The *Saccharomyces cerevisiae* (*S. cerevisiae*) yeast strain VL6-48N was used for TAR cloning [63]. The genotype of this strain is: *MATα, his3-Δ200, trp1-Δ1, ura3-Δ1, lys2, ade2–101, met14,* cir°. Yeast was grown in YPD media supplemented with adenine (400 μg/ml) at 30°C. Yeast transformed with the YCp/BAC was grown in synthetic dropout (SD) media (Sigma) lacking histidine (−HIS) or without uracil (−URA) and supplemented with adenine. *E.coli* strain TOP10 (Invitrogen) was used for the propagation of all BAC-cloned TAR plasmids and assembled genomes. Cultures were grown overnight at 37°C in 2YT media containing chloramphenicol (25 μg/ml).

### Plasmids

The EBV Akata-BAC^LHF^ was a generous gift from Dr. Lindsey Hutt-Fletcher (LSU Health Sciences Center, Shreveport, LA). This is an Akata genome cloned into the BAC vector. Yeast plasmids pCC1BAC-ura3 and pCC1BAC-his3 [64] have been previously reported. pEZT-BM was a gift from Ryan Hibbs (Addgene plasmid # 74099) {University of Texas Southwestern Medical Center, Dallas, TX [65].

### Oligonucleotides and Synthetic DNA

Oligonucleotides were synthesized by Invitrogen or IDT, and gBlocks were generated by IDT. TrueGuide gRNAs were designed using CHOPCHOP software [66] and synthesized by Invitrogen.

### Whole Plasmid Sequencing

All TAR plasmids were sequenced using the Nanopore sequencing method by Psomagen, Poochon Scientific or Plasmidsaurus. Whole genome sequencing of EBV^YA^ WT 3.11.1 and WT 10.1.1 was carried out by the MGH CCIB DNA Core. The DNA was digested with I-SceI first, purified using AMPure XP magnetic beads and the library preparation done using end ligation-based methods.

### EBV^YA^ Genome Analysis using Hybrid-Capture Next-Generation Sequencing (NGS)

To determine the sequence content of the assembled yeast genomes, we performed NGS combined with a hybrid-capture target enrichment strategy as previously described ([67]). This approach was developed by the Xian laboratory to perform high-throughput whole-genome sequencing of both major strains of EBV (Type I NC_007605 and Type II NC_009334). EBV-specific probes were synthesized as single-stranded DNA probes comprising 120-base sequences tiling the entire EBV genome (Integrated DNA Technologies, Coralville, IA). Probes targeting both non-repetitive and repetitive sequences were designed to generate complete viral genome sequences. In brief, after adapter ligation and PCR, the sequencing libraries from each EBV^YA^ genome underwent hybrid-capture enrichment using this panel of EBV-specific probes. Following enrichment, the libraries were amplified, purified, and sequenced on the MiSeq instrument (Reagent Kit v2, 300-cycles, Illumina Inc). FASTQ files were aligned to three reference genomes NC_007605, NC_009334 and human genome build hg19 to generate BAM files that were visualized in IGV ([68]). Sequencing depth was also calculated at each viral genomic coordinate.

### Synthesis of vectors for transformation-associated recombination (TAR) cloning

Vectors were PCR amplified using pCC1BAC-his3 or pCC1BAC-ura3 as templates with KOD Xtreme Hot Start polymerase (Millipore-Sigma). This plasmid contains a bacterial artificial chromosome (BAC) and yeast centromeric plasmid (YCp) sequence for replication in *E.coli* and yeast, respectively. The construction primers add an I-SceI restriction enzyme site, flanked by 40 bp of EBV homology, to each end of the vector backbone (Table S1). KOD PCR products were digested with DpnI (New England Biolabs [NEB]) prior to gel isolation and transformation.

### Lithium acetate yeast transformation

Lithium acetate transformation of yeast was adapted from Bartel and Fields [69] and Desai et al. [70] and modified by Oldfield et al. [30], as follows, EBV Akata DNA, pCC1 vector DNA, and 150 μg of carrier DNA (Takara) were mixed in an Eppendorf tube. Yeast cells (100 μl) were added to the DNA, followed by 650 μl of a 50% PEG/10X TE/1M lithium acetate mixture (8:1:1 v/v/v). This transformation mix was incubated at 30°C for 30 min, followed by the addition of 70 μl of DMSO and heat shock at 42°C for 20 min. Transformants were plated on the appropriate selective SD media plates and incubated at 30°C.

### TAR cloning, screening, and processing of EBV DNA cloned fragments

Three different EBV templates were used for TAR cloning: Akata, Akata-BX1 and Akata-BAC^LHF^. EBV DNA from lytically induced cells was purified using the Hirt extraction method, whereas EBV Akata-BAC^LHF^ DNA was isolated using PureLink HiPure Plasmid Filter DNA Purification Kit (Invitrogen) that uses anion-exchange resin for high purity DNA preparations. We used both the Midi and Maxi formats. Generally, for the Midiprep, we used 100 ml of culture in 2YT broth. The volumes of the resuspension, lysis and precipitation solutions were increased (12 ml per 50 ml culture). In addition, after the neutralization step, we clarified the mixture by centrifugation prior to loading it onto the column. The DNA was eluted from the columns using elution buffer that was at 65°C. The template DNAs were prepared for TAR cloning by shearing (pipetting 10-20x) or by restriction enzyme digestion. Restriction enzymes that cut on either side of the TAR fragments were used to create free DNA ends for TAR cloning. BAC template DNA (600 ng) or infected cell DNA (5 μg) was co-transformed with 20 to 230 ng of the appropriate vector into competent yeast cells. Transformants were patched on SD minus HIS plates. Once grown, yeast was picked into 25 μl of 25 mM NaOH and incubated at 95°C for 30 minutes to lyse the cells. Q5 High-Fidelity DNA polymerase (NEB) PCR on DNA from lysed yeast was used to confirm the correct junction of the vector and EBV fragment (Table S2). Each individual yeast was screened by PCR assays using Vector F-TAR Det R (e.g. for TAR 2: J2-3 Det R) and Vector R-TAR Det F (e.g. for TAR 2: J2-3 Det F). For TAR 11 we used Vector R507-TAR Det F (Table S2).

Zymolase DNA preparations were made of positive clones from liquid cultures grown in SD minus HIS as described by Oldfield et aI. [30]. Isolated DNA was then electroporated into competent *E. coli* TOP10 cells (Invitrogen). Single *E. coli* colonies were screened again for vector-EBV junction sequences using Phire Hot Start II DNA polymerase (Invitrogen). DNA was purified from positive transformants using the PureLink HiPure Plasmid Filter DNA Purification Kit. Each TAR clone was designated with the strain and fragment number (e.g. EBV TAR 4; Fig. 1). Each TAR clone was checked by restriction enzyme digestion (Fig. 2) and sequenced using Nanopore methods (Table 1 and Fig. S1) three-four times using different companies (Psomagen, Poochon Scientific and Plasmidsaurus).

**Fig. 2.**
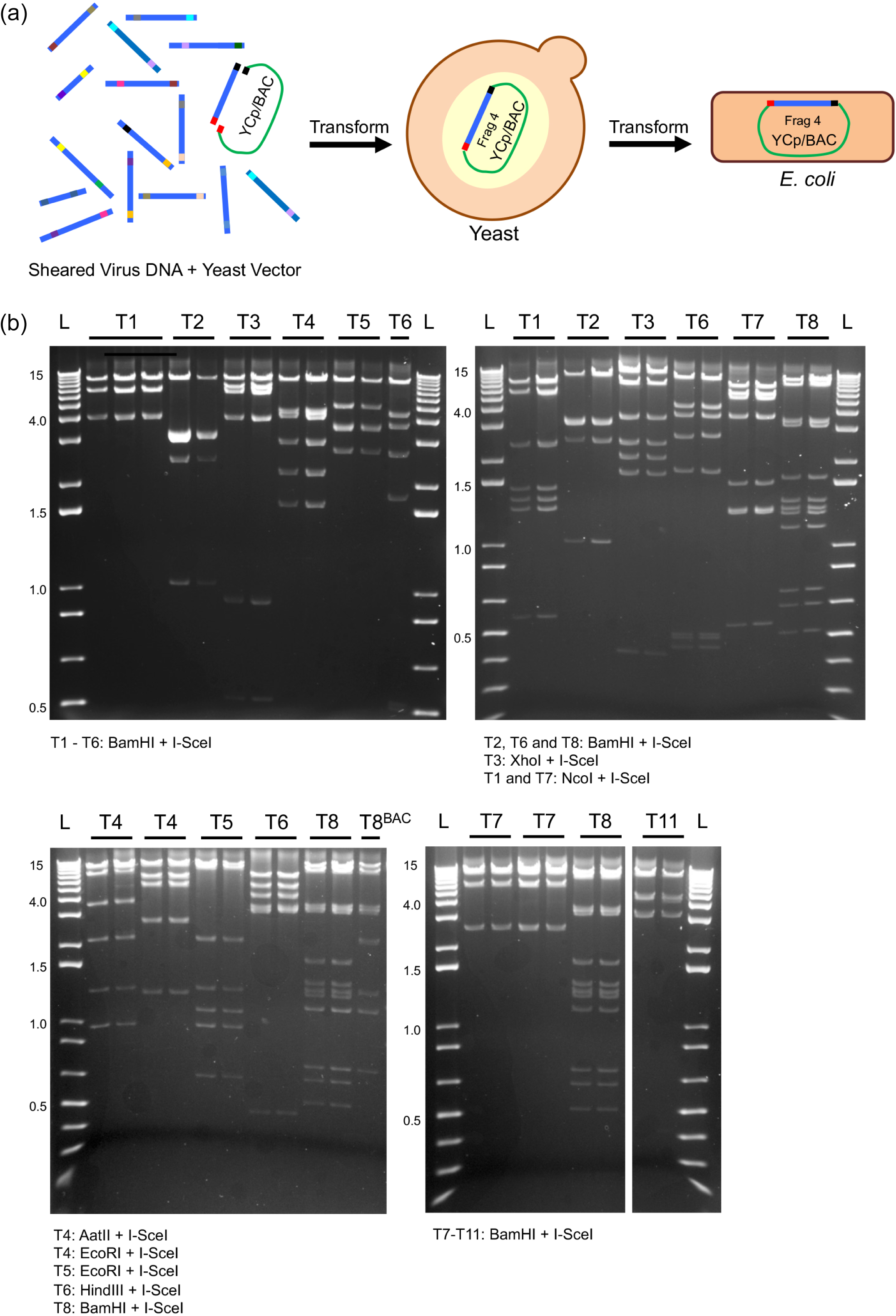
TAR cloning of EBV fragments TAR 1 to TAR 11. (a) Schematic representation of the general methodology used to clone the eleven TAR fragments. A YCp/BAC vector encoding a histidine (HIS) selection marker was utilized to isolate each TAR fragment. Sheared or restriction enzyme-digested EBV Akata DNA served as templates. (b) TAR plasmids were propagated in TOP10 *E. coli*, and purified DNA was analyzed by restriction enzyme digestion. All the TAR plasmids shown in this figure are derived from Akata-BX1 with the exception of TAR 8 for which we show the clone derived from the Akata-BAC^LHF^ genome. DNA ladders are in the first and last lanes, with sizes in kilobases (kb) indicated on the left. Multiple clonal isolates of the TAR plasmids were analyzed. TAR 9 and TAR 10 were derived later and were analyzed by long read sequencing.

**Table 1.**
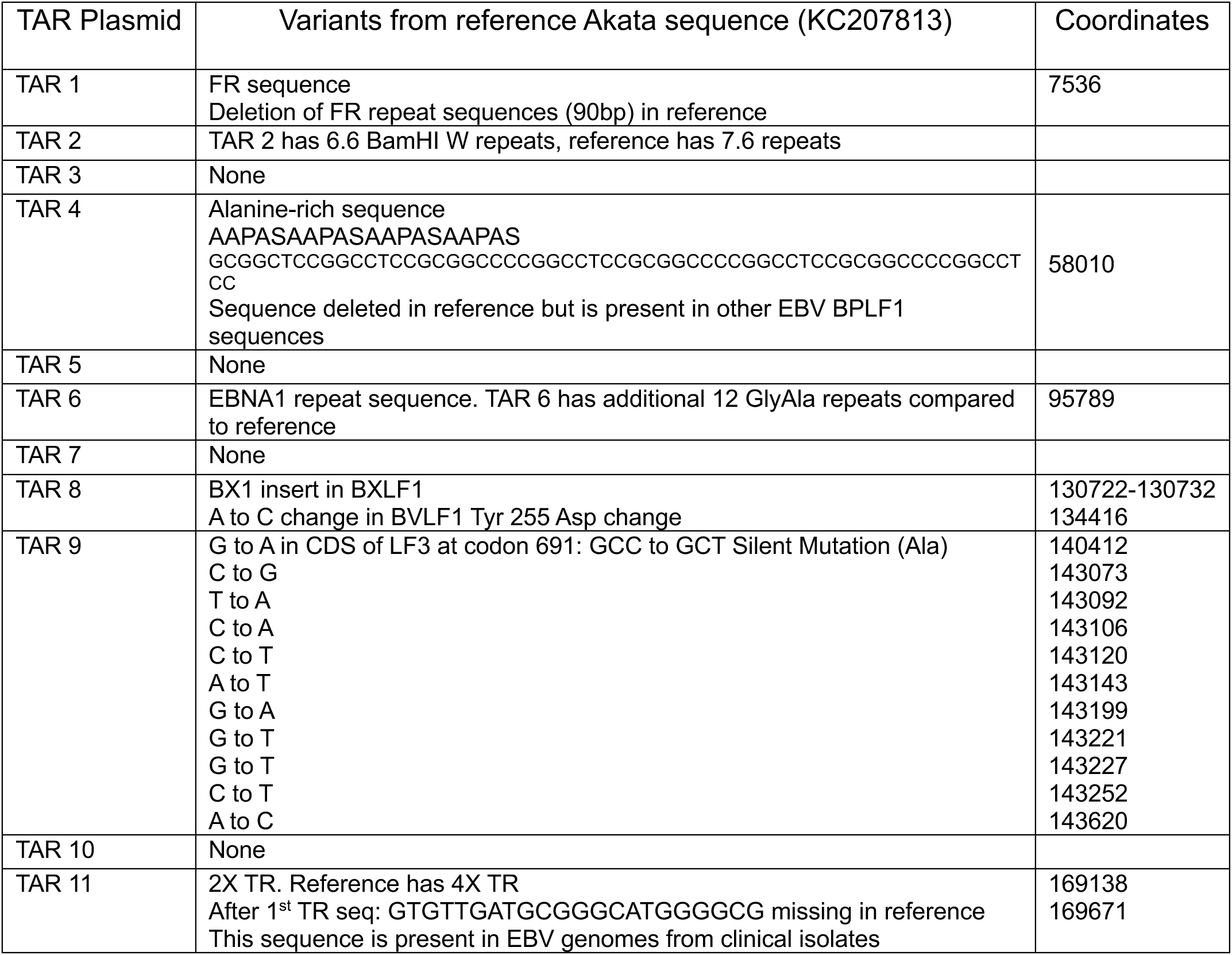
Akata-BX1 TAR plasmid sequence variants from reference.

### Engineering Deletions in BFRF3 and BZLF1

A deletion in the *BFRF3* open reading frame (ORF) was engineered using CRISPR/Cas9-guided digestion followed by TAR cloning in yeast. Due to the overlapping genes at the *BFRF3* genomic loci, a complete deletion of the ORF could not be generated. Thus, we designed a deletion encompassing codon 42 to 123 out of a total of 176 amino acids. We used TAR 3, which contains *BFRF3* ORF, as the template (Fig. 1). A TrueGuide Synthetic gRNA (Table S3) was designed and purchased from Invitrogen. This gRNA and Cas9 enzyme (NEB) were used *in vitro* to cleave TAR 3 at amino acid 66 position, using the manufacturer’s protocol. Cleavage was confirmed by agarose gel electrophoresis. Both the cleaved TAR 3 and a gBlock containing the desired *BFRF3* deletion were transformed into yeast VL6-48N for TAR cloning. The gBlock encoded 90 bp of homology on either side of the deletion. The resulting yeast colonies were patched and screened for the *BFRF3* deletion using Q5 PCR assays (Table S3). Using this method, we could generate the desired mutants with high efficiency (8 out of 11 patches analyzed). Zymolase DNA preparations were carried out for positive clonal isolates and electroporated into TOP10 cells. These were analyzed, and purified DNA preparations were isolated as described above for the TAR clones. The purified TAR 3-ΔBFRF3 plasmid was sent for Nanopore whole-plasmid sequencing to validate the deletion and the TAR 3 backbone sequence.

Similarly, a deletion for *Zta* (*BZLF1*) was designed using the same methods. A complete deletion of the *Zta* ORF (only the ATG and TAG were retained) was generated because there are no overlapping genes at this genomic locus. We used TAR 6 which contains the *Zta* ORF, as the template (Fig. 1). A TrueGuide synthetic gRNA was designed (Table S3), purchased from Invitrogen, and used together with Cas9 enzyme to cleave TAR 6 at codon 110 of *BZLF1*. Again, yeast cells were transformed with the cut TAR 6 and a gBlock which has 90 bp homology arms on either side of the *Zta* deletion. Confirmation of the deletion and validation of the mutant TAR 6 were performed using the same methods described for the *BFRF3* deletion.

### Genome Assembly

The final EBV^YA^ genome has the YCp/BAC vector at the ends of the virus genome, between TAR 11 (after the TR region) and TAR 1. Because of this configuration, we determined that half-genome sub-assemblies should be constructed first, followed by a full assembly from the two halves. The two sub-assemblies generated include TAR 1–6 (T1–6) and TAR 7–11 (T7–11), which were subsequently joined in a full-genome assembly to generate TAR 1–11 (T1–11) (Fig. 3). Because all individual TAR plasmids are cloned into pCC1BAC-his3, it was necessary to implement a counter-selection strategy during these assembly steps.

**Fig. 3.**
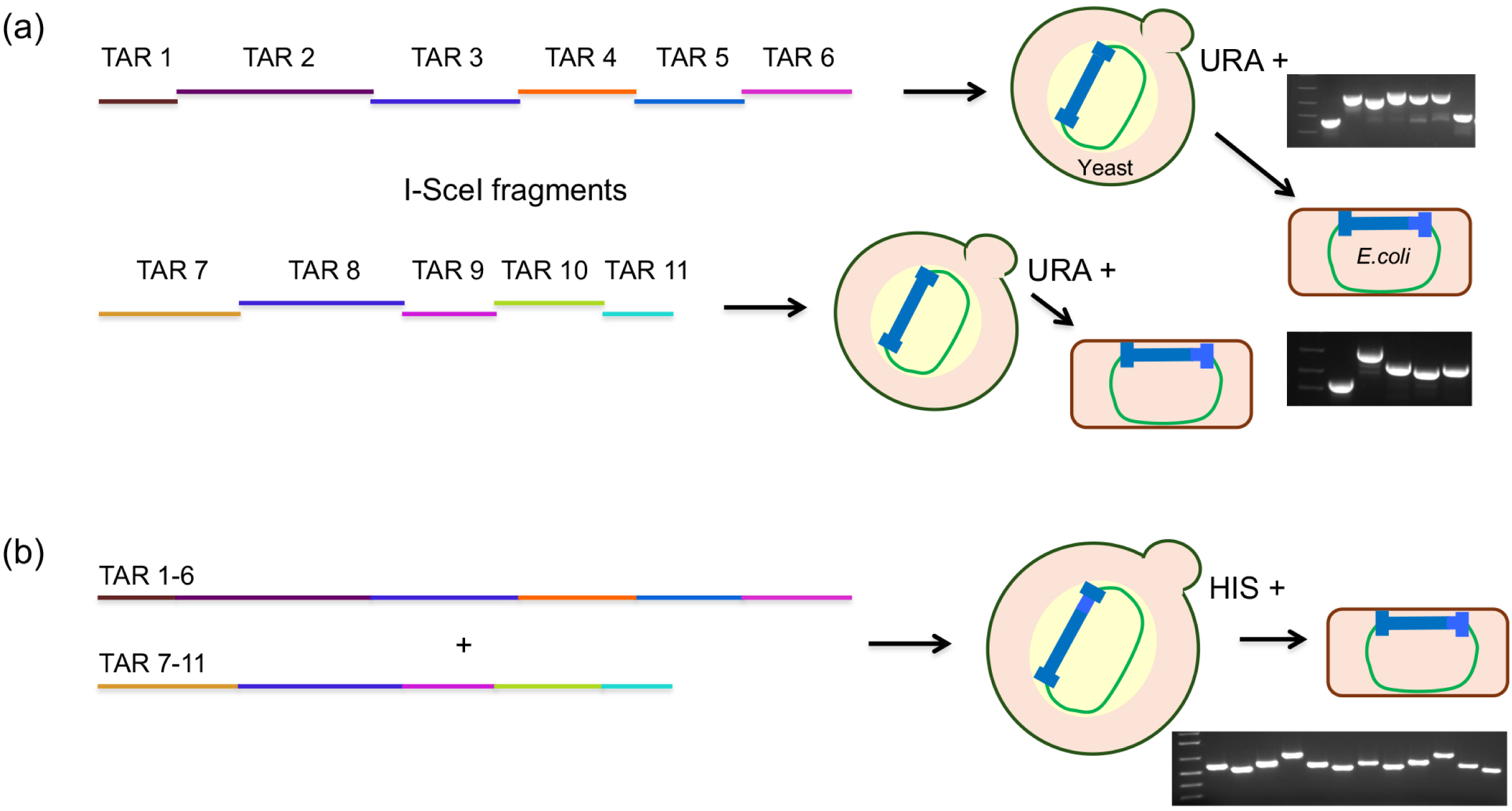
Modular assembly of TAR fragments into a complete genome. (a) Three-step assembly strategy for the EBV Akata-BX1 genome. First, individual TAR fragments were released from the YCp/BAC histidine (HIS) vector by I-SceI digestion. Sub-genomes TAR 1–6 and 7–11 were independently assembled by TAR cloning in yeast using a YCp/BAC vector carrying an uracil (URA) selection marker. Assembled half-genomes were screened using junction-detection PCR. (b) The TAR 1–6 and TAR 7–11 genomes were transferred into *E. coli*, and the isolated DNA was digested with I-SceI and co-transformed into yeast with the YCp/BAC HIS vector. Complete genomes were identified by screening yeast cells for all DNA junctions. Positive clones were transferred into *E. coli* and validated by restriction enzyme digestion (Fig. S3 and S4).

Prior to assembly of the TAR fragments, 10 μg of each TAR plasmid was digested with I-SceI (NEB) to release the EBV fragments from the vector backbone. The I-SceI-digested DNAs were visualized via agarose gel electrophoresis to quantify the relative amounts of DNA based on fluorescence intensity. For each half-genome assembly, 1 μg of each TAR plasmid fragment was combined with 100–200 ng of pCC1BAC-ura vector for yeast transformation. The yeast transformants selected on SD minus URA plates were patched out for analysis using Q5 polymerase PCR assays. Both single-junction and nested PCR assays were performed to identify transformants that successfully assembled a half-genome (Tables S2 and S4 and Fig. S2a-b). Single-colony isolates were derived from the positive yeast patches and grown in SD minus URA medium overnight. Zymolyase DNA preparations were performed, and the isolated DNA was electroporated into competent *E. coli* TOP10 cells. Single bacterial colonies were cultured overnight in 2XYT medium containing chloramphenicol and then screened again for the presence of each junction using Phire polymerase. A positive clonal isolate was grown, and the plasmid DNA was purified using the PureLink HiPure Plasmid Filter DNA Purification Kit. The purified half-genome sub-assembly was subsequently validated via restriction enzyme digestion analysis (Fig. S3a-c).

After verification, a whole genome assembly was conducted using the T1–6 and T7–11 sub-assemblies. I-SceI digestion was performed on both sub-assemblies, and the digests were visualized using agarose gel electrophoresis to ensure equimolar amounts of DNA were used. The half-genomes were co-transformed into competent yeast VL6-48N cells along with the pCC1BAC-his3 vector containing 40 bp sequences homologous to either end of the EBV genome. The transformed yeast was plated onto SD minus HIS plates this time for counter-selection, and the resulting yeast colonies were patched and screened for all junctions using Q5 PCR assays (Fig. S2a-b).

### Transfer of complete EBV genomes from yeast into *E. coli*

The assembled EBV genomes (both half and whole) were handled with extreme care to avoid shear forces generated by vortexing and pipetting. We used only genomic (large bore) tips for handling these genomes. Single-colony isolates were derived from the positive yeast patches and grown in SD minus HIS medium overnight. We adapted a yeast zymolase DNA extraction method described by Boeke and colleagues [71]. Ten ml of overnight yeast culture grown in SD-HIS at 30°C was processed and the cells pelleted and resuspended in 500 μl of Solution I (8 ml 1M Sorbitol and 2 ml 0.1M EDTA pH 7.5). Cells were again pelleted in a microcentrifuge and the pellet resuspended in 400 μl of Solution II (8 ml 1M Sorbitol and 2 ml 0.1M EDTA pH 8 and 100 μl of β-mercaptoethanol). 100 μl of zymolase 20T, 10 mg/ml (Zymo Research) was added to the cells, inverted gently to mix and incubated at 37°C for 30 minutes (inverting gently every 15 minutes). The spheroplasts were pelleted in a centrifuge and then gently dislodged from the bottom with 400 μl of TE buffer. To this, 90 μl of Solution III (0.9 ml of 0.5M EDTA pH 8, 1.2 ml Tris pH 8 and 0.6 ml 10% SDS) was added and the suspension, incubated at 65°C for 30 minutes (inverting gently after 15 minutes). Then 80 μl of 5M potassium acetate was added, the suspension inverted gently to mix and incubate on ice for 30 minutes. The mixture was centrifuged for 15 minutes at 4°C. The supernatant was transferred to a new Eppendorf tube and the DNA precipitated by adding 1 ml of cold ethanol. The DNA was pelleted and the pellet was washed with 70% ethanol, air dried and resuspended in 200-400 μl of TE buffer. We stop here in the Boeke protocol. The subsequent RNase and phenol/chloroform extraction steps in the Boeke protocol were not carried out. Competent *E. coli* TOP10 cells were electroporated with 20 μl of the DNA preparation. Single *E.coli* colonies were grown overnight in 2XYT medium containing chloramphenicol and then screened for the presence of the full assembly (J1 to J10) using Phire polymerase PCR with nested detection primers (Table S4 and Fig. S2b). EBV DNA from positive clones was isolated using the PureLink HiPure Plasmid Filter DNA Midiprep Kit (Invitrogen) and confirmed by restriction enzyme digestion analysis (Fig. S4a-b) and whole genome sequencing (Fig. 4 and Fig. S5a-b).

**Fig. 4.**
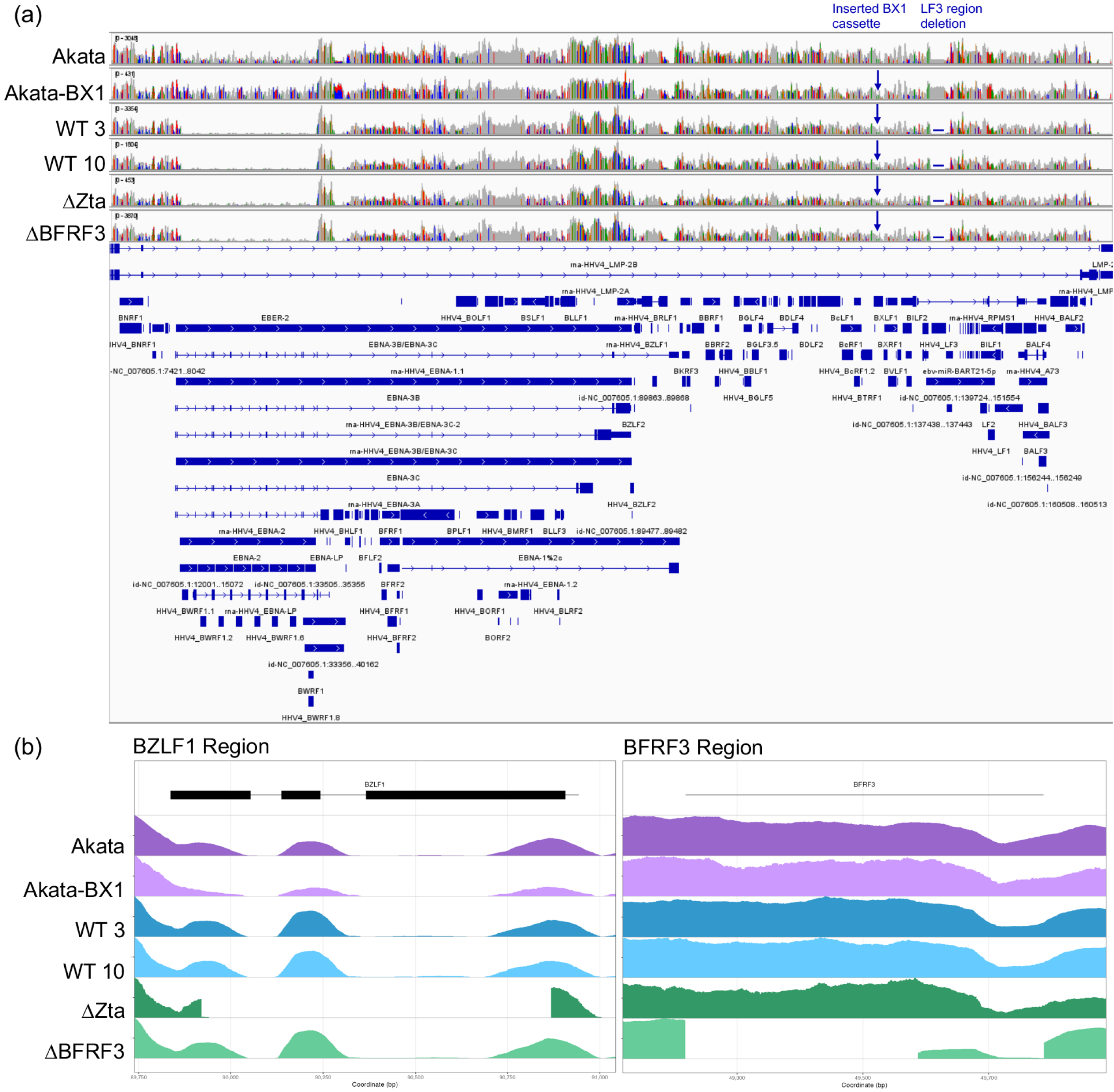
Hybrid-capture sequencing of wild-type (WT) EBV^YA^ genomes and the ΔZta and ΔBFRF3 mutants using MiSeq. (a) Genome-wide coverage tracks from six samples—Akata, Akata-BX1, and the yeast-assembled EBV genomes WT3, WT10, ΔZta, and ΔBFRF3—showed broad and continuous coverage across the EBV genome. The yeast-assembled samples showed reduced coverage in the BamHI-W region compared with Akata and Akata-BX1. Note that Akata and Akata-BX1 were sequenced from genomic DNA in a different sequencing run than the EBV^YA^ samples. The recombinant viruses contain the same cassette insertion in the BXLF1 region as Akata-BX1. The yeast assembled genomes also show a deletion in the LF3 region. (b) Zoomed-in coverage plots of the *BZLF1* and *BFRF3* regions showed reduced or no coverage across the corresponding deleted regions in the ΔZta and ΔBFRF3 genomes, respectively, consistent with the engineered deletions.

### Transfection of EBV genomes into 293 cells

The assembled EBV genomes were transfected into HEK-293 cells (6 x 10^6^ cells in a 12-well tray) using 5-10 μg of PureLink purified DNA and 8 μl X-tremeGENE HP DNA Transfection Reagent (Roche), according to the manufacturer’s instructions. After 2–3 days, GFP-positive cells were visualized, and the cells were trypsinized. The cells were then transferred to 100 mm dishes for selection with Geneticin (G418; Invitrogen) starting at an initial concentration of 0.5 mg/ml. After 3 days, the drug concentration was lowered to 0.25 mg/ml. The dishes were incubated until individual GFP-positive colonies were detected. Single colonies were isolated using cloning cylinders and expanded in complete DMEM containing 0.25 mg/ml Geneticin. The clonal cell lines were expanded in 24-well plates, subsequently scaled to 6-well plates, and finally transferred to T75 flasks in the continuous presence of Geneticin (0.5 mg/ml).

### Recombinant Baculoviruses

For generation of recombinant baculoviruses expressing Zta (*BZLF1*), Rta (*BRLF1*), glycoprotein B [gB] (*BALF4*), and small capsid protein (*BFRF3*), we used the plasmid pEZT-BM [65] (Fig. S6a), which is engineered for expression of cloned genes in mammalian cells. The BacMam (BM) plasmid utilizes the HCMV immediate-early (IE) promoter to drive expression of the cloned genes. It also contains an mEGFP cassette under the control of the baculovirus p10 promoter for expression in insect cells. Thus, viruses generated in insect cells can be visualized by mEGFP fluorescence, which also serves as a marker for virus titer in insect cell cultures (Fig. S6b). *BZLF1* and *BFRF3* were cloned as XhoI-KpnI fragments; *BRLF1* was cloned as an NheI-KpnI PCR amplified fragment; and *BALF4* was cloned as a NotI-KpnI fragment into pEZT-BM (Table S3 lists the PCR primers used for cloning). A cDNA copy of *BZLF1* (IDT) served as the template for PCR amplification. TAR 3 and TAR 10 DNA was used as templates for *BRFR3* and *BALF4* respectively and BX1-Akata genomic DNA was used as the template for *BRLF1*. The BZLF1 ORF was also cloned into the baculovirus vector pFB1 mCherry. This vector as described in Desai et al. generates a C-terminal fusion of mCherry to the protein [72]. This plasmid was then used to amplify *BZLF1*-mCherry and cloned as an XhoI-KpnI fragment into pEZT-BM. All ORFs were PCR-amplified using Q5 High-Fidelity DNA Polymerase, using the manufacturer’s protocol. Cloned ORFs were sequenced to verify the absence of PCR-generated mutations (Fig. S7a-d). Baculoviruses were generated using the Bac-to-Bac Baculovirus Expression System (Invitrogen). The protocols for transfection, production, and amplification of baculoviruses are documented in detail by Perkins et al. [73] and Okoye et al. [74].

### Virus Production from HEK-293 Cell lines

HEK-293-EBV^YA^ cells (4.5 x 10^6^ cells) were seeded into 100 mm tissue culture dishes in DMEM supplemented with 10% FBS. Upon reaching 90% confluency, lytic replication was induced by transducing cells with recombinant baculoviruses expressing Zta, Rta and gB. The baculovirus transduction mixture consisted of 300 μl of each virus and 600 μl of PBS. Generally, our baculovirus stocks are 10^8^ plaque forming units/ml as shown by Okoye et al. [74]. This mixture was incubated with the cells for 8 h at 37°C, after which time the inoculum was removed and replaced with complete DMEM containing 1 mM sodium butyrate. At 96 h post-transduction, the supernatant was harvested and clarified by centrifugation at 1493 x *g* for 15 min using a benchtop centrifuge. The clarified viral supernatant was concentrated in 5 ml tubes by centrifugation at 20,900 x *g* for 1 h. The supernatant was removed, and the viral pellet was resuspended in 100 μl of PBS overnight at 4°C. This concentrated viral stock (final volume with residual media was 150 μl) was subsequently used for primary B-cell infections and Raji cell titrations.

### Raji GFP Titration

Raji cells were utilized to qualitatively analyze viral production efficiency from the producer cells. Raji cells (2 x 10^6^ cells) were resuspended in 100 μl of RPMI media containing 20 ng/ml 12-O-tetradecanoylphorbol-13-acetate (TPA) and 1 mM sodium butyrate (designated as RTN media). The cells were incubated with either concentrated virus (50 μl out of 150 μl stock) or in some cases unconcentrated virus supernatant (500 μl out of 4.5 ml stock) in a 24-well tissue culture plate. The plates were incubated at 37°C for 8 h or overnight, after which, the viral inoculum was replaced with fresh RTN media. Cells were imaged for GFP fluorescence using a Bio-Rad ZOE Fluorescent Cell Imaging System.

### B cell Purification and Infection

Human blood samples were obtained with informed consent under an IRB approved protocol (IRB NA 00028682). Human bone marrow mononuclear cells (MNCs) were purified by Ficoll gradient density sedimentation. B cells were further purified using two different methods. The first approach used CD19 MicroBeads (Miltenyi Biotec) for positive selection of CD19+ B cells, using the manufacturer’s protocol. Subsequently, B cells were isolated via negative selection using the RosetteSep Human B Cell Enrichment Cocktail (STEMCELL Technologies). This latter method was found to be more effective at purifying "untouched" primary B cells. Following a second wash with PBS post-Ficoll separation, cells were resuspended in a mixture of 500 μl of PBS, 600 μl of red blood cells, and 250 μl of RosetteSep cocktail. This mixture was incubated at room temperature for 20 minutes with gentle agitation every 5 min. RPMI media (15 ml) was then added to the mixture, which was layered onto a Ficoll gradient and processed as described above. Purified B cells were either cryopreserved in 10% DMSO or adjusted to a concentration of 2-5 million cells/200 μl of RPMI media for subsequent infection with concentrated EBV (50 μl of stock). Note that when negative selection was used, no calcineurin inhibitor was used. Infected cells were monitored for several days to visualize the formation of LCLs.

### Reagent Sharing

We are committed to sharing all reagents which include DNA plasmids, genome assemblies, recombinant viruses, cell lines and protocols generated in this study. Our history of sharing reagents is evident from the numerous laboratories to which we have distributed HSV-1 recombinants—especially K26GFP—as well as complementing cell lines for capsid and tegument mutants.

## RESULTS

### Molecular design of the modular assembly of the EBV Akata-BX1 genome

The design of the cloning of EBV fragments was based on the Akata genome which is resident in a BL cell line. Our choice to use the Akata strain was because many of our laboratory investigations are based on Akata specifically Akata-BX1 which is an engineered recombinant made by Lindsey Hutt-Fletcher [52]. This clone has a GFP reporter/neomycin selection cassette inserted in the *BXLF1* (*TK*) gene (Fig. S1b) which allows one to visualize cells harboring the genome using fluorescence microscopy and to maintain the latent genome using G418 selection. The Akata published sequence from Flemington et al. [40] was used as the reference sequence (GenBank: KC207813.1) (Fig. 1). This genome of 171.3 kb consists of unique sequences as well as several repetitive elements. These include the terminal repeats (TR), the large internal repeats (IR1) also known as the BamH1W repeats (BamW), as well as origins of latent and lytic DNA replication that contain repetitive sequences (Fig. 1). The genome was broken up into 11 parts/fragments. The borders of those fragments were adjusted to locate repeat sequences internally and not at the ends (Fig. 1). Each fragment overlaps the neighboring fragment by 79 unique base pairs on each side (Table S1). The vector sequence contains both BAC and YCp sequences at the same site for growth in *E. coli* and *S. cerevisiae*. This vector sequence is designed so that it spans the ends of the Akata genome. It is inserted after the TR region. It can be excised by cleavage with I-SceI (Fig. 1). The complete assembled EBV genome was designated EBV^YA^ (EBV yeast assembled).

### Transformation-associated recombination (TAR) cloning of eleven fragments that span the EBV Akata-BX1 genome

The 11 overlapping fragments were isolated by TAR cloning, which uses the natural propensity of *S. cerevisiae* to undergo homologous recombination for the repair of double-strand DNA (dsDNA) breaks to clone large regions of DNA [60]. A TAR cloning vector with BAC and YCp sequences was generated by PCR amplification (Table S1). The 3′ and 5′ ends of each vector contain 40 bp of homology to a targeted, unique EBV region, as well as sites for the I-SceI restriction enzyme, which does not cleave the EBV genome, to allow for release of the cloned fragments. Each of the 11 fragments was cloned in independent reactions by co-transforming the TAR cloning vector into yeast cells with sheared EBV genome template DNA, either from Akata, Akata-BX1 (purified from lytically induced cells) or Akata-BAC^LHF^ DNA purified from *E. coli* (Fig. 2a). These transformants were selected by growth on HIS selection plates. DNA from positive TAR clones, identified by PCR with detection primers (Table S2) across the junction of the vector and virus fragment, were transformed into *E. coli* and were further confirmed by restriction enzyme digestion analysis (Fig. 2b) as well as Nanopore sequencing (Table 1). Positive clones were designated with the TAR and clone number (e.g., EBV TAR 4 for fragment 4). We derived all eleven TAR fragments from Akata-BX1 genomic DNA. These were the 11 we used for whole genome assembly. We did derive TAR 2, 3, 4, 8, 9 and 11 from Akata-BAC^LHF^ DNA and TAR 9 and 10 from Akata genomic DNA. The Akata-BAC^LHF^ clones were primarily a back-up but also TAR 8 is required if we need to assemble a genome with an intact *BXLF1* (TK) gene. For most TAR clones we used sheared template DNA (pipetting 10-20x). In some cases, we used restriction enzymes that cut on either side of the TAR fragment to generate free ends for TAR cloning. These were for BX1: TAR 2 (NdeI/HindIII), TAR 3 (SnaBI/EcoRI), TAR 8 (PsiI), TAR 10 (AatII/AclI) and TAR 11 (ScaI/EcoRI).

Nanopore sequencing of the TAR clones from BX1 revealed variants from the reference sequence (KC207183) [40]. These are shown in Table 1. Most of the variants are in the repeat regions. For example, there is a deletion in the FR region in both the Flemington and Kanda Akata FR18 sequence and the Kanda Akata FR23 sequence has three more additional FR sequences (Fig. S1a) [55]. There are 6.6 BamW repeats in TAR 2 and 7.6 in the reference. The BamW is 3 kb in size and 0.6 is the partial 1.8 kb unit. There is an alanine-rich sequence in tegument protein BPLF1 that is missing in the reference sequence but is present in other EBV BPLF1 sequences. The TAR 6 EBNA1 region has

12 additional GlyAla repeats compared to the reference. There are also only 2 TR sequences compared to the 4 in the reference. In addition, there is a sequence after the first TR that is missing in the reference sequence but is present in sequences of EBV clinical isolates. There is one amino acid change in BVLF1 (TAR 8) and silent mutations in LF3. There are also several variants in the TAR 9 sequence just outside of the LF3 sequence. The BX1 cassette is in TAR 8 and consists of the CMV promoter driving the expression of GFP and an HSV-1 TK promoter driving expression of the Neomycin gene. There are also SV40 and BGH poly(A) sequences. The insert is after codon 464 of BXLF1 (Fig. S1b).

The stability of the TAR plasmids has been examined in *E.coli* and yeast following serial passaging in those hosts [30]. We do not see any instability of the EBV TAR plasmids following yeast replication and isolation from bacterial hosts. We have amplified these TAR plasmids many times without any detrimental effects on their structures. This includes the repeat regions in the different TAR plasmids such the BamW, FR and the other internal repeat regions. This is based on restriction enzyme profiling, PCR detection assays, whole plasmid sequencing using Nanopore sequencing and Sanger sequencing.

### Assembly of a complete EBV genome in yeast from 11 overlapping fragments

In our EBV genome design, the pCC1 vector (YCp/BAC) is located at the ends of the genome rather than internally, as was designed for HSV-1 [30]. The vector also is not in the TR region but outside of that sequence. Because of this configuration, we determined that half-genome sub-assemblies should be constructed first, followed by a full assembly from the two halves. The two sub-assemblies generated include TAR 1–6 (T1–6) and TAR 7–11 (T7–11), which were subsequently joined in a full-genome assembly to generate TAR 1–11 (T1–11) (Fig. 1 and Fig. 3). All EBV fragments were released from the vector sequence by I-SceI digestion, co-transformed, and assembled by TAR in *S. cerevisiae*, which produces a scarless junction without any non-native sequence or I-SceI restriction sites (Fig. 3a). The resulting transformants were selected for on uracil-deficient plates and screened by PCR to confirm the presence of junctions between adjacent fragments, indicating successful assembly of the EBV half-genome (Fig. 3, Fig. S2a and Table S2). The confirmed half-genomes were transferred to *E. coli* to generate DNA preparations for restriction enzyme analysis. We generally used BamHI, EcoRI and HindIII to analyze the integrity of the half-genomes and to eliminate any genomes with deletions in the repeat elements (Fig. S3a-c) [9]. The number of BamW repeat units which are 3 kb in size do vary in a few different assemblies. This was evident in TAR 1-6 (Fig. S3a, lane 3 of first panel, WT 9.1) which is missing all the 3 kb fragments. The BamHI repeats are also located in the largest EcoRI and HindIII fragments and differences in the mobility of these fragments relative to the second largest fragment can reveal changes in the number of repeats (Fig. S3a-b and S4a-b). Genomes that may have been generated by non-homologous end joining and contain deletions can also be visualized by these analyses (Fig. S3a lane 2 first panel, WT 6.1). The family of repeat (FR) region within oriP was also analyzed by a PCR assay that can detect any deletions in this element (Fig. S2c). Occasionally we have seen deletions in this region during genome assembly. Hence, the PCR screen is done on all genome assemblies (half and whole). We also confirmed the FR is similar in TAR 1, Akata and Akata-BX1 genomic DNA and the Akata-BAC^LHF^ (Fig. S2d). Even though the PCR assay is difficult in this region due to GC content and secondary structure [55] our current assay is able to identify deletions that could potentially impact genome maintenance. Once the half-genomes were confirmed, we assembled the whole genome. Again, the now larger fragments (TAR 1-6 and TAR 7-11) were released by I-SceI digestion and these pieces together with the vector were co-transformed into yeast cells (Fig. 3b). This time the colonies were selected for on histidine-deficient media plates and colony patches were screened by PCR for all junctions (J1 to J10) spanning the EBV genome (Fig. S2a-b). Positive clonal isolates were again transferred into *E. coli* and analyzed by restriction enzyme digestion (Fig. S4a-b) and PCR assays. During sub-genome and whole genome assembly we do observe missing DNA sequences as discussed above. Using the junction PCR detection methods we preferentially isolate genomes that assemble by homology mediated recombination versus non-homologuos end-joining. Nevertheless, we have seen changes in the number of the BamW repeats and occasionally the FR repeat deletion. It is likely as more fragments and thus free DNA ends are transformed into yeast there is an increase in the recombination activity, hence the appearance of more errors in the genome assembly.

### EBV^YA^ whole genome sequencing using NGS

Whole viral genome sequencing, both short-read (Fig. 4, wild-type, ΔZta and ΔBFRF3) and long-read (Fig. S5a-b, wild-type), demonstrated complete EBV genomes in all yeast-assembled EBV genomes apart from a region overlying LF3 and the IR4 PstI repeat region (NC_007605.1:140,851-143,339), which was not recovered at all from the enriched short-read sequencing libraries, and was only recovered infrequently from the long-read sequencing libraries. All EBV^YA^ genomes contained the BX1 cassette at position NC_007605.1:131,174-131,179 leading to a deletion of these 6 bases of the EBV genome. Lastly, based on relative sequencing depth observed over IR1 BamHI W Repeat when compared to other cell lines, such as Akata, the number of W repeats may be reduced, though the exact number of repeats cannot be determined (Fig. 4a). Regarding specific mutant constructs, sequencing confirmed deletions of the following gene regions: BZLF1 and BFRF3 (Fig. 4b). Neither the short-read or the long-read sequencing methods were able to resolve accurately the number of BamW repeats.

Based on the sequencing data, we again compared different TAR 1–11 assemblies to the TAR 2 plasmid using restriction enzyme digestion (Fig. 5a). The TAR 1–11 DNAs were digested with EcoRI and HindIII, which give rise to a fragment that spans the IR1 repeat region. TAR 2, which contains 6.6 BamW repeats, was digested with I-SceI to release the fragment from the YCp/BAC. The TAR 2 fragment has a Nanopore-sequenced size of 21,845 bp which includes the 6.6 BamW repeats and is used here as a molecular ruler (white dot). The EcoRI–HindIII fragment spanning the IR1 (yellow dot) is predicted to be 25,259 bp. Because the seventh repeat is absent in TAR 2, we subtracted 3073 bp from the predicted Akata EcoR1-HindIII fragment size (28,332 bp). The difference between the EcoRI-HindIII fragment and TAR 2 fragment is 3,414 bp. On the gel we observed that the two fragments migrated with similar mobility in the TAE buffer system used, even though there is a 3.4 kb (∼1 BamW repeat) difference (Fig. 5a) Hence, we conclude that the TAR 1–11 genomes analyzed here have a minimum of 5.6 BamW repeats.

**Fig. 5.**
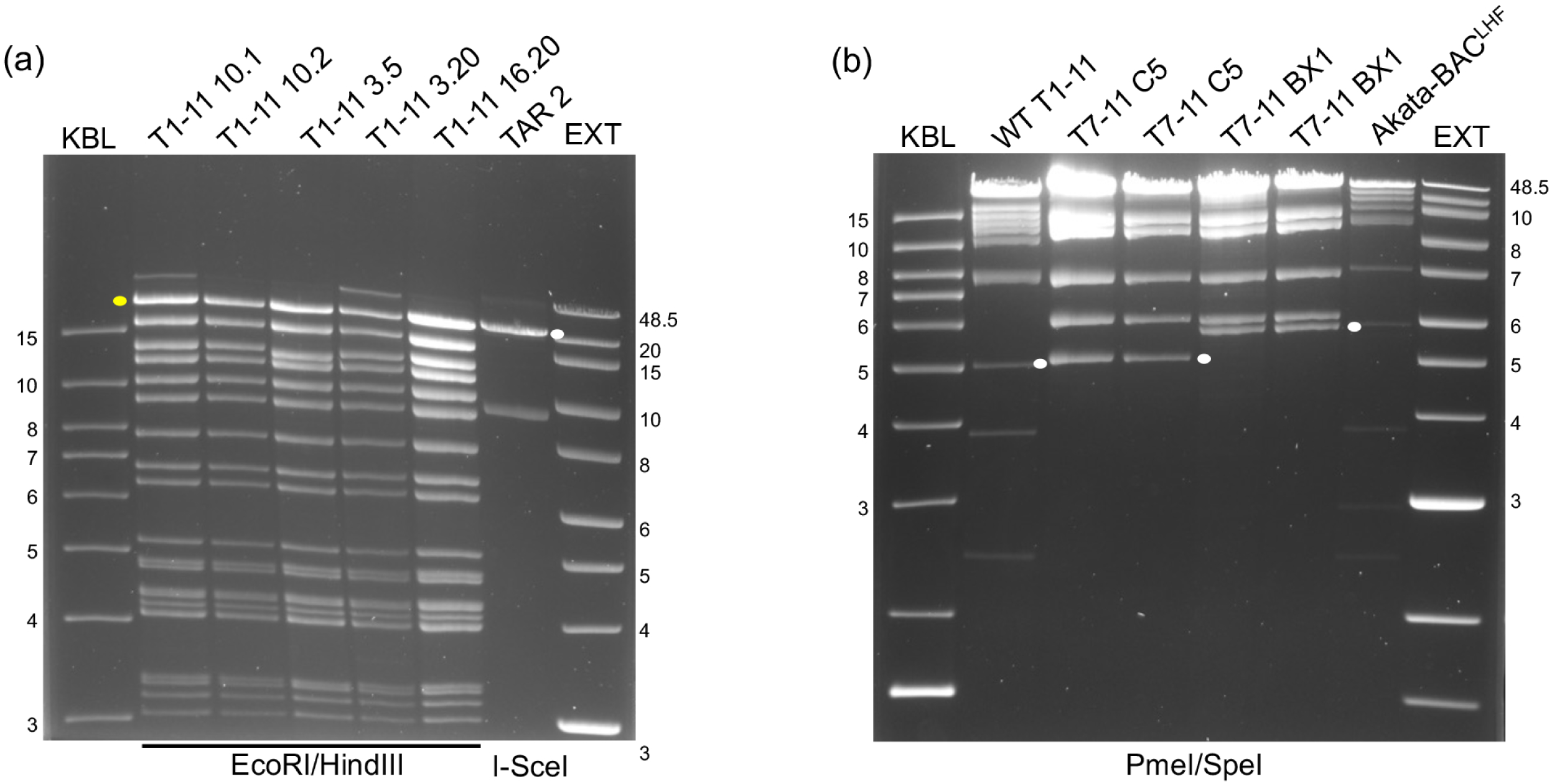
Gel electrophoresis analysis of BamW repeat region and IR4 repeat region. (a) EBV^YA^ TAR 1-11 genomes were double-digested with EcoRI and HindIII restriction enzymes and the fragments resolved on a 0.9% TAE gel (35V). The TAR 2 plasmid was digested with I-SceI. The TAR 2 fragment contains 6.6 BamW repeats and has a predicted size of 21,845 bp, white dot. The EcoRI-HindIII fragment spanning IR1 has a predicted size of 25,259 bp (a 3,414 bp predicted difference), yellow dot. (b) Wild-type (WT) TAR 1-11 and TAR 7-11 genomes (clone 5 and BX1) were double-digested with PmeI and SpeI and the restriction enzyme fragments analyzed by gel electrophoresis as described above. The fragments of interest are indicated by white dots. Two different preparations of the TAR 7-11 clones were analyzed. DNA ladders are shown in lane KBL (1kb + ladder) and EXT (1 kb extend ladder). The sizes (kb) of the standards are indicated.

There was also a loss of sequence content in the IR4 repeat region in all the yeast-assembled clones, indicative of a deletion in the sequenced region. Because the deletion is present in all assemblies, it likely occurred at the stage of assembly of TAR 7–11 (clone 5), which contains the IR4. To resolve this, we looked again at our assemblies and checked both TAR 1–11 and two different TAR 7–11 assemblies: clones 5 and BX1 36.2 (Fig. 5b). We used TAR 7–11 clone 5 for all the whole-genome assemblies in this paper. There is a SpeI–PmeI fragment that spans this region (white dot), and its calculated size is 5,875 bp. When we examine this fragment in these assemblies, the fragment is shorter in TAR 7–11 clone 5 (approximately 5 kb) compared to TAR 7–11 BX1 36.2 fragment, which has a mobility around the 6 kb marker (Fig. 5b). The TAR 7–11 clone 5 band is similar in size to the fragment in TAR 1–11 (WT). The predicted deletion, based on this gel analysis, is approximately 700-800 bp in size. The extremely high GC content of the LF3 gene is likely the DNA deleted as judged by partial Sanger sequencing in this region. The oriLyt is intact based on the hybrid-capture sequencing.

### Generation of HEK-293 cells as the host for infectious virus production

To make cell lines for virus production, we used HEK-293 cells. Due to the presence of a neomycin resistance gene in the BX1 cassette, parental HEK-293 cells were used instead of the large T antigen-transformed line. DNA was transfected into cells using Xtreme-Gen HP transfection reagent which gave us the highest transfection efficiency of this large genome. Transfected cells were followed by GFP fluorescence and cells were selected in the presence of G418 (Fig. 6a). Colonies that were GFP+ and G418 resistant were harvested using cloning cylinders and amplified. These cell lines are always grown with media containing 0.5 mg/ml G418. We monitor GFP expression routinely and have not observed any loss of GFP signal on passage.

**Fig. 6.**
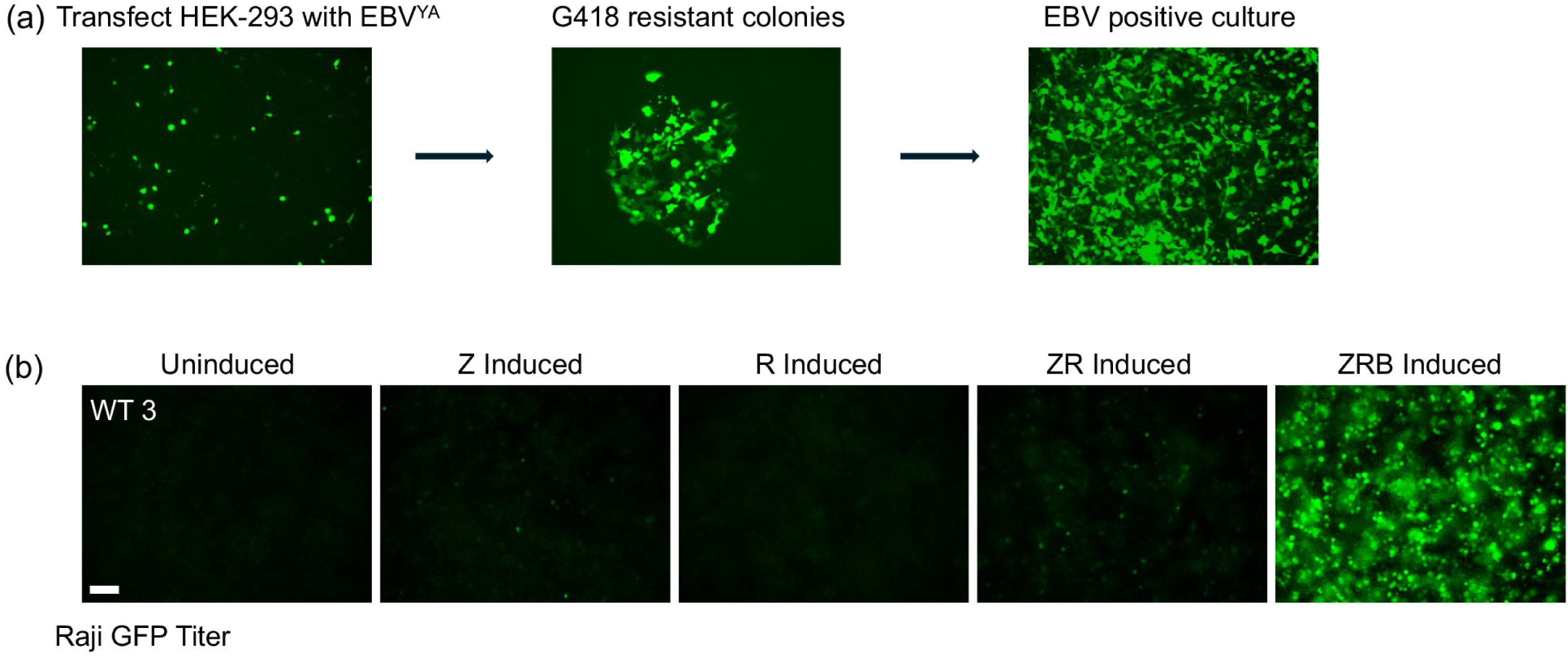
Generation and analysis of EBV virus producer cell lines. (a) The yeast assembled (YA) EBV genomes were transfected into HEK-293 cells. Clonal lines were isolated following G418 selection and GFP fluorescence visualization, then expanded prior to evaluating virus production. (b) Clonal lines harboring wild-type (WT) genomes were induced into lytic replication by transduction with baculoviruses expressing Zta (Z), Rta (R), and glycoprotein B (B). Virus yields, quantified by Raji GFP titers, peaked following co-transduction with all three Zta, Rta, and gB-expressing baculoviruses. These results were observed repeatedly with multiple virus producer cell lines. Scale bar = 100 μm.

### Baculovirus-mediated transduction of Zta, Rta and gB of HEK-293 cell lines for infectious virus production

Because the EBV genome is dormant in the HEK-293 cell lines, lytic inducers such as Zta and Rta provided *in trans* are required to turn on virus production as has been shown previously [75]. Following the experiments by the Kenney and Delecluse laboratories, the inclusion of gB (BALF4) is also required [76, 77]. Recent studies have shown the expression of BALF4 (gB), BVLF1 and BKRF4 enhanced virus production after lytic induction [78]. Hence, GFP+, and G418 -resistant cell clones were first screened for lytic induction by providing Zta, Rta and gB *in trans*. Previously, this has been done by transfecting plasmids expressing the three proteins or combinations of them [9, 13, 59, 75-79]. Based on our history of baculovirus gene expression, we used recombinant baculoviruses engineered for mammalian gene expression (BacMam) [80]. The ORFs of *Zta*, *Rta* and *BALF4* were expressed from the CMV IE promoter (Fig. S6a and S7a-d). This baculovirus-based transduction method has been shown to effectively induce KSHV virus production through KSHV RTA expression [81]. Furthermore, with baculoviruses we can achieve a high level of transduction of HEK-293 cells as judged using a fluorescent reporter (Zta-mCherry) expressing baculovirus (Fig. S8). Virus secreted into the supernatants of induced cells was concentrated by centrifugation and used to infect Raji cells. We initially determined that the expression of all three genes was the best combination for virus production as judged by the number of GFP+ Raji cells (Fig. 6b). We then tested several protocols to optimize lytic induction and virus production. We varied the time of infection from 2 h to 8 h (Fig. 7a) and the time of harvest from 24 h to 96 h (Fig. 7b). After multiple inductions tests, we used an 8 h infection period for the baculoviruses and the culture supernatant was harvested at 96 h post-infection (Fig. 7). During this induction period, the HEK-293 cells were monitored using the fluorescence microscope to discover which clones displayed an increase in GFP signal, indicative of lytic induction. We choose cell lines that demonstrated robust lytic induction as judged by the increased GFP signal. The culture supernatants were harvested from these clones and concentrated by centrifugation. This virus was used to infect Raji cells to determine the levels of virus production. Many cell lines gave high numbers of GFP-positive Raji cells, indicating the yeast assembled virus is biologically infectious and can produce virus capable of infecting lymphocytes. We have attempted different methods to quantitate virus titers, such as, qPCR and flow cytometry without much success in the rigor of the data. However, we now present quantitation of Raji GFP Units (RGU) that was determined by counting GFP positive cells following serial dilution (Fig. S9). The typical titer averaged approximately 2 x 10^5^ RGU/ml from a high producer cell line. For most of our infections (Raji and B cells) we used 50 μl, that is, one third of the concentrated stock.

**Fig. 7.**
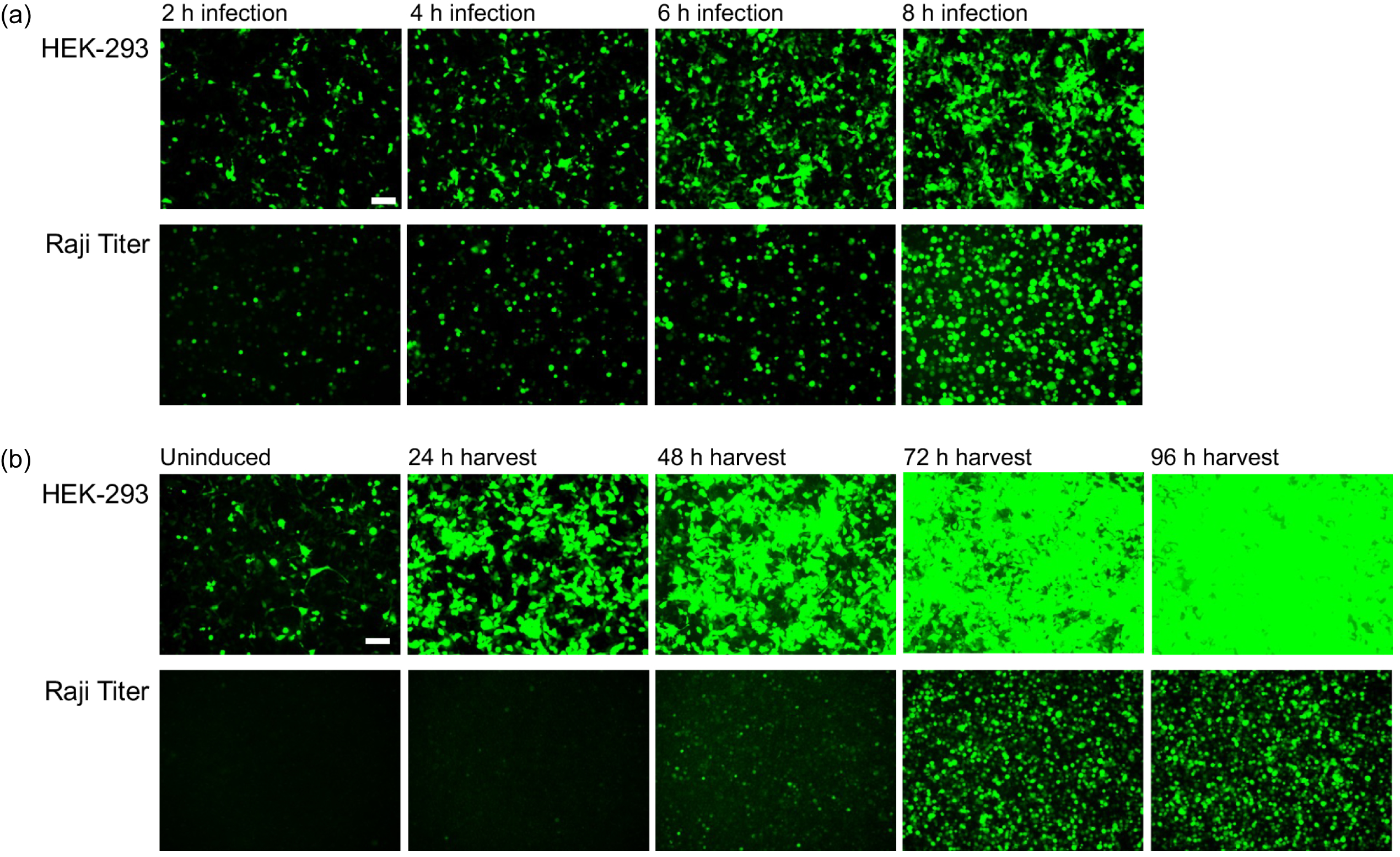
EBV^YA^ WT virus producer cell lines: kinetics of virus transduction and virus production. (a) HEK-293 lines were infected with baculoviruses for the indicated time points. Supernatant-harvested virus was concentrated and used to infect Raji cells. An 8-hour baculovirus infection of HEK-293 cells gave the highest virus yield as judged by the GFP signal in Raji cells. (b) Supernatants from induced cells (8-hour baculovirus infection) were harvested at different time points post-infection and concentrated by centrifugation. Raji cells were infected to quantify infectious virus yields based on the number of GFP-positive cells. Similar results were observed with different HEK-293 EBV^YA^ cell lines. Scale bar = 100 μm.

### Yeast assembled EBV can immortalize resting B cells into lymphoblastoid cells

The Raji titration assay demonstrated that the HEK-293 cell lines were producing high levels of infectious virus. The next step was to examine whether this virus could transform primary B cells. Rather than infect PBMCs, we purified B cells and infected them with concentrated EBV^YA^ virus. Initially, CD19+ cells were purified from bone marrow donors using positive selection. These cells were infected with EBV^YA^ and cyclosporin A was included in the culture media to inhibit cytotoxic T cells. Subsequently, we used a negative selection approach, with RosetteSep B cell enrichment technology. This time cyclosporin A was omitted during culture of the infected cells. In both cases, the cells were cultured and followed for the appearance of (LCLs) (Fig. 8a). Initially, we observed clusters of a few cells that were not strongly GFP+ (Fig. 8a), but as the cells grew into larger clusters the GFP signal became much stronger (Fig. 8b). We were able to derive LCLs using B cells purified by both methods; however, the B cell enrichment method gave the better infection and more efficient method of producing LCLs (Fig. 8c). Mock infected B cells did not result in any outgrowth of LCLs.

**Fig. 8.**
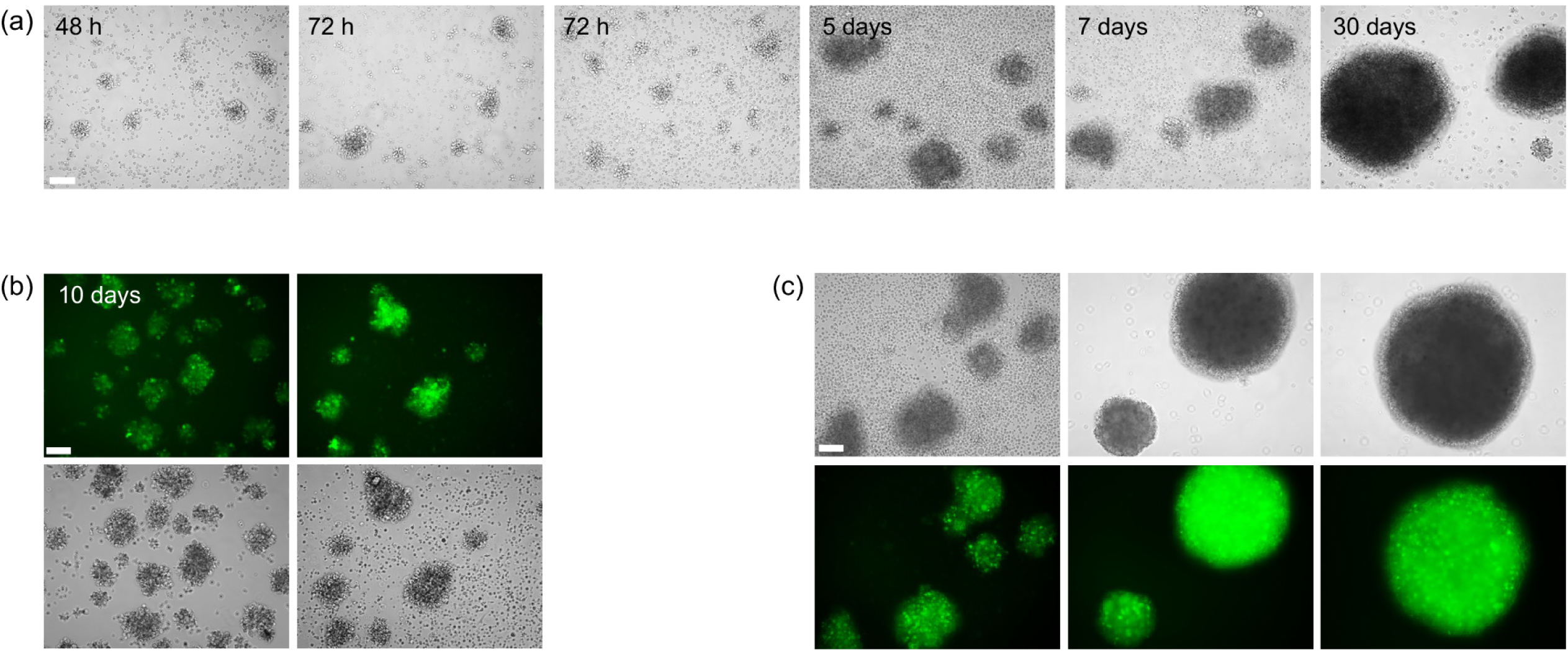
Immortalization of B cells by EBV^YA^virus. Primary B cells were infected with different wild-type virus preparations and monitored for transformation into lymphoblastoid cell lines (LCLs). (a) Time-course of B-cell infection progressing to LCL formation at 30 days post-infection. (b) Phase-contrast and fluorescence microscopy revealed small clusters of infected, GFP-positive cells at 10 days post-infection. (c) Large, strongly green, fluorescent LCL clusters observed at 1–2 months post-infection. Similar results were seen during transformation of B cells with different virus preparations. Scale bars = 100 μm.

### Engineering null mutations of *BZLF1* and *BFRF3* in EBV^YA^ and complementation *in trans* of these lethal phenotypes using baculovirus transduction

To test the utility of the EBV genome assembly method, two deletion mutations were engineered. One was in *Zta* and the other was in the gene encoding the small capsid protein (SCP), *BFRF3*. Many investigations have examined deletions of *Zta* and have shown that this is an essential gene of EBV [13, 75]. For *BFRF3*, the effect of a null mutation on virus replication was unknown. The HSV-1 SCP is not required for virus growth in cell culture but does have a role in virus replication in mouse ganglia [82]. Using the recombinant baculovirus expression system in insect cells, we showed that the SCP of EBV is required for capsid assembly in Sf21 insect cells [83]. For *Zta*, we made a null mutation (Fig. 9a), while for *BFRF3*, due to overlapping open reading frames, a deletion of the internal codons 42 to 123 was made (Fig. 10a). Both mutations were generated using a CRISPR Cas9 approach, which we have also applied successfully to HSV-1 engineering [30]. This method is highly efficient and yields the desired changes in the TAR plasmid. Modification was done on the individual TAR fragment, TAR 6 for *Zta* and TAR 3 for *BFRF3*. *In vitro* Cas9 cleavage with a targeted gRNA was used to introduce linear ends within the gene of interest sequence. The “cut” TAR fragment together with a gene block that encoded the required deletion and 90 bp of homology on either side of the deletion was transformed back into yeast for repair and introduction of the desired change. TAR plasmids were screened for the deletion using PCR assays and then, the whole plasmid was sequenced. Again, when these engineered plasmids were sequenced, we did not observe any instability or deletion of the repeat regions and sequences. The engineered parts were introduced into the TAR 1 to 6 assembly reaction and subsequently into the TAR 1-11 whole genome assembly.

**Fig. 9.**
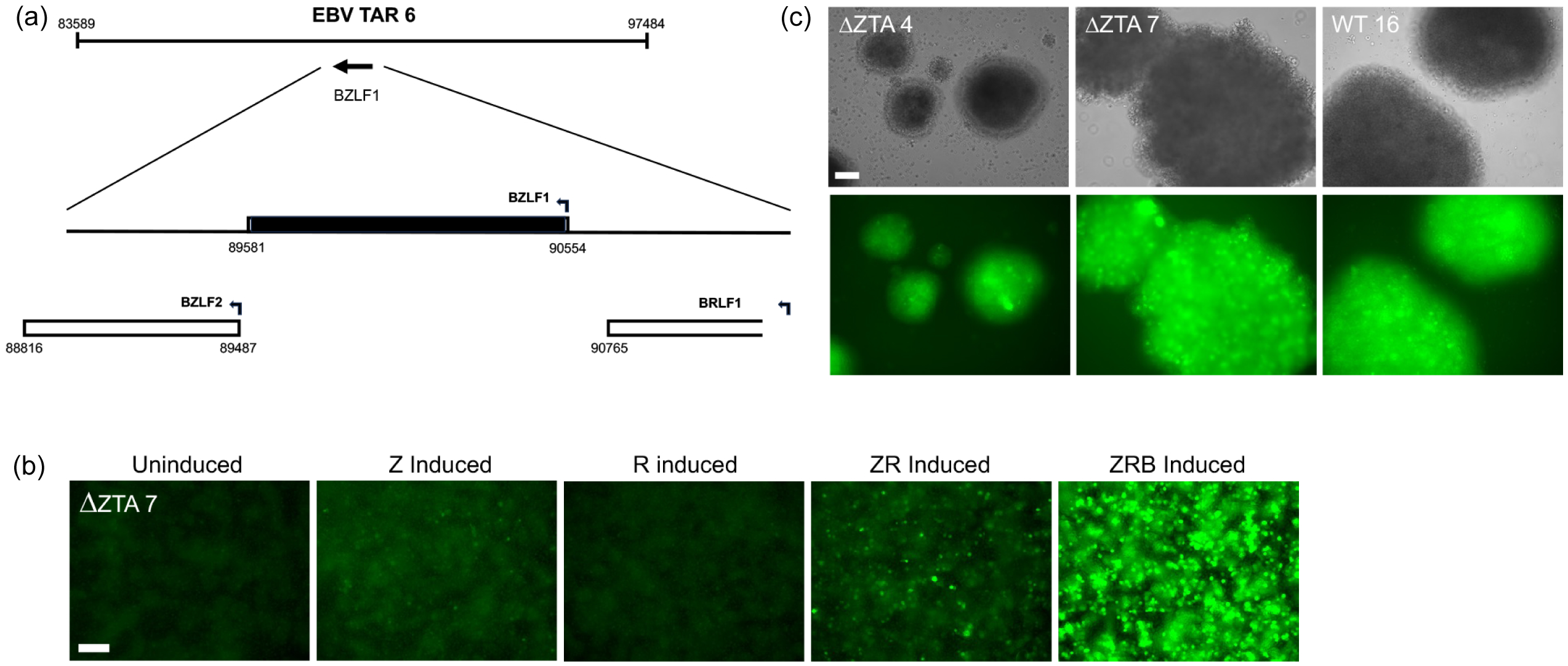
Analysis of an EBV^YA^ genome encoding a deletion of *BZLF1*. (a) A deletion within the *BZLF1* gene (black shading) was engineered into TAR 6, which was sequence-validated and used for whole-genome assembly. EBV Akata genome coordinates and transcriptional directions (arrows) are indicated. (b) HEK-293 virus producer cell lines were established following transfection with EBV^YA^ ΔZta. Representative clone 7 was induced with baculoviruses expressing lytic activators Zta (Z), and Rta (R) and glycoprotein B (B). Infectious virus, quantified by Raji GFP titer, was detected only upon induction with Zta, Zta+Rta, or Zta+Rta+gB. (c) B cells were transformed into LCLs using virus derived from wild-type or ΔZta producer lines (clones 4 and 7). These results were seen in multiple experiments with the different ΔZta mutant cell lines. Scale bar = 100μm.

**Fig. 10.**
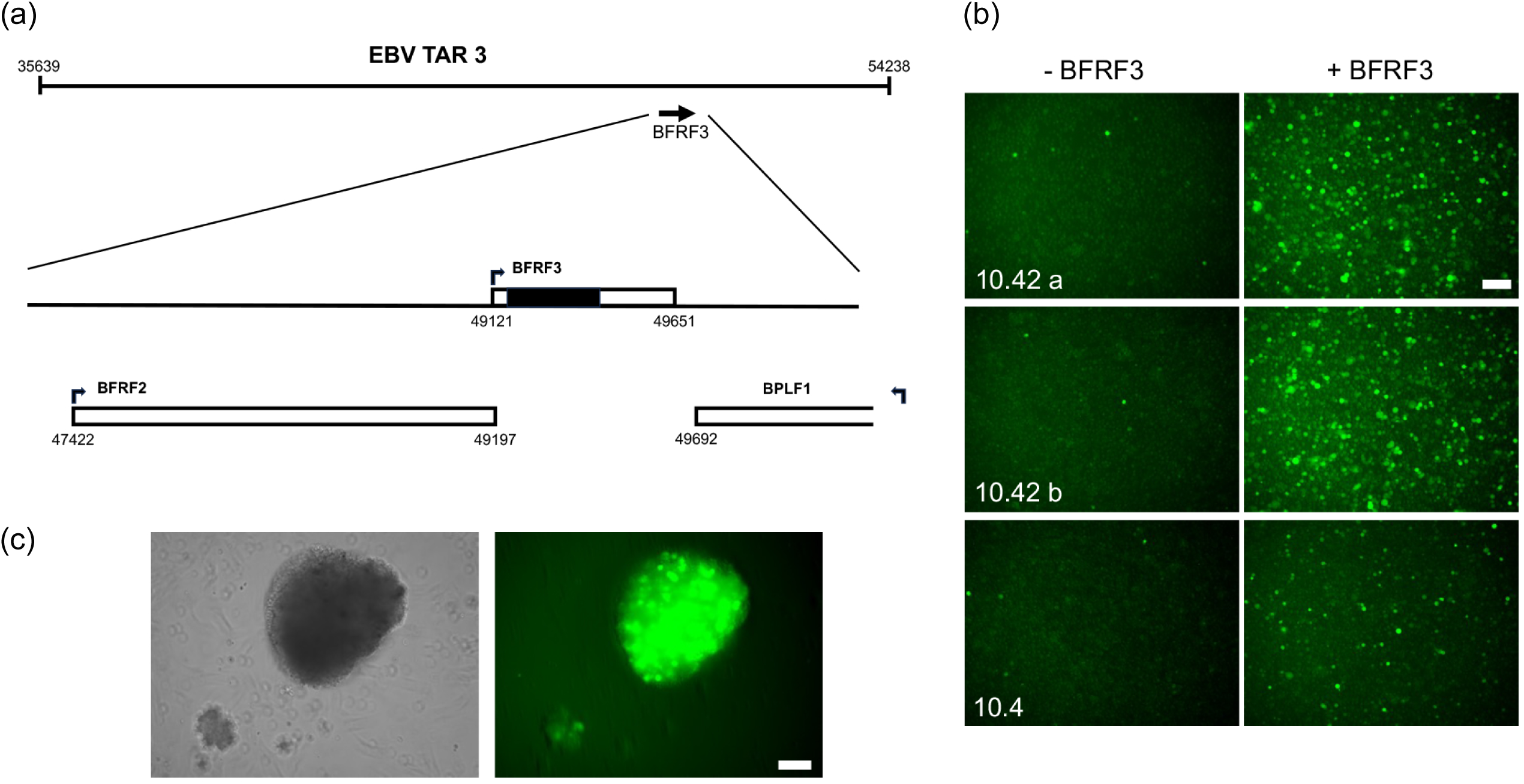
The small capsid protein of EBV (BFRF3) is essential for virion formation. (a) A deletion within the *BFRF3* open reading frame (ORF; black shading) was engineered into TAR 3 and incorporated into the EBV genome assembly in yeast. EBV Akata genome coordinates and transcriptional directions (arrows) are indicated. (b) HEK-293 producer cell lines were established for the EBV^YA^ ΔBFRF3 genome. These cells failed to produce virus when induced with baculoviruses expressing Zta, Rta, and gB (–BFRF3), as measured by Raji GFP titers. Complementation with a BFRF3-expressing baculovirus (+BFRF3) rescued virion production, as confirmed by GFP fluorescence in Raji cells. Consistent results were observed across multiple independent HEK-293 clones. (c) Rescued virus particles successfully infected and transformed primary B cells into LCLs. Scale bar = 100μm.

The ΔZta and ΔBFRF3 genomes were transfected into HEK-293 cells to generate virus producer cell lines. Several clonal lines were derived that showed a significant increase in the expression of GFP following lytic induction using baculovirus transduction. For the ΔZta cell lines, virus production, as judged by Raji titration, was observed only when the baculovirus expressing Zta was included in the lytic induction (Fig. 9b). The virus complemented by Zta *in trans* could transform B cells into LCLs (Fig. 9c).

For the ΔBFRF3 cell lines, virus was not detected when the cells were induced for lytic gene expression, even though GFP expression increased. However, when BFRF3 was provided *in trans* using baculovirus gene expression, virus was detected using the Raji assay (Fig. 10b). This last result demonstrated the importance of BFRF3 for EBV replication and the ability to complement lethal mutant virus production using a baculovirus expression method. This virus could also transform B cells into LCLs (Fig. 10c).

## DISCUSSION

Previous studies from our group demonstrated the ability to build a complete herpesvirus genome using multiple fragments that overlap with each other in yeast [30, 31]. The premise for this is based on the assembly of large bacterial genomes using cloned fragments. TAR cloning was a significant method development that allowed one to precisely design the cloning of large fragments [61, 84-86]. This study demonstrates that an EBV genome assembled in yeast is biologically functional, producing virus capable of transforming B cells. More recently, TAR cloning has been used to clone out whole herpesvirus genomes. This method, which is termed STAR (single step TAR) cloning has been used to isolate two strains of Rat CMV as well as new CMV strains of mouse and derive cloned genomes from KSHV cell lines, BCBL-1 and HBL-6 [87, 88]. In addition, TAR cloning was used to assemble genomes of HSV-1 strain H129-Syn that can be used for neuronal tracing [89-91]. These yeast-based synthetic biology methods have been leveraged to clone many different virus genomes and chromosomes [84, 91-97].

The use of BAC-cloning technology [98] was a transformational change for herpesvirus researchers after Messerle et al. demonstrated the cloning of the mouse cytomegalovirus genome in a BAC [1]. Since then, many herpesvirus genomes have been cloned using BAC plasmids [3, 9, 11, 12, 14, 20, 25, 27, 28, 99] and once in *E. coli*, can be modified by recombineering methods [15, 100-102]. This advance enabled the introduction of reverse genetics to many herpesviruses that previously could not be manipulated easily [3, 9, 27, 28]. However, some problems working with these cloned genomes have been reported [55, 103]. In addition, the allelic exchange methods used tend to expose the whole genome, albeit for a short duration, to very active recombinases that could potentially generate unwanted changes. These issues can greatly complicate the generation of targeted mutations and interpretation of phenotypes. The synthetic genomics method has potentially significant advantages over the BAC-based system. The synthetic genome assembly method divides the genome into multiple parts like an automobile assembly line. The different parts can be changed separately, prior to assembly of the final genome, and any problem part can be modified in isolation and then put back into the assembly line. Unwanted mutations can also occur in yeast cells when the fragments are assembled into genomes by non-homologous end-joining recombination events. We are able to mitigate these events by screening using junction PCR detection assays, thus excluding such unwanted changes at the junctions from further analysis. While the individual TAR plasmids are generally stable during recombineering, the larger genome builds may inherently incorporate errors because of the increased number of DNA free ends in yeast. Another factor that affects the use of herpesvirus-BACs for reverse genetics is the time needed to make changes in the genome. In the synthetic assembly line, genome-wide changes can be made rapidly because we can manipulate any designated number of the parts individually or in parallel to create the desired alterations and then, combine these altered parts with the rest of the parts in the assembly line to obtain the modified complete genome. This assembly approach also provides a more stable system for engineering the genome. Repeat sequences and DNA origins of replication that can cause instability of the genome are ‘stored’ in the host cells separately from each other and are only present at the final step when the complete genome is assembled.

Some of the observations made using Akata-BX1 as a template for EBV genome assembly were the variations in the BamHI-W (BamW) repeat region (IR1). We cloned TAR 2 with different numbers of BamW repeats using Akata-BX1 as template. The TAR 2 fragment that we used for EBV genome assembly had a copy number of 6.6 repeats. Nanopore sequencing of different DNA preparations of the TAR 2 plasmid clone consistently showed 6.6 W repeats indicating this was a stable clone. The Akata genome sequence contains 7 BamW repeats plus a partial 8 repeat, therefore, 7.6 [40]. However, occasionally during EBV sub-genome assembly, we did see a change in this region, as judged by restriction enzyme digestion of the assemblies. We eliminated the genomes with lower numbers of repeats. Previous studies have shown using an EBV episome plasmid that 5 BamW repeats are required for maximal transformation of B cells. They did not test the efficiency of transformation with 4 BamW repeats. Transformation is reduced with lower numbers of the repeats (3 BamW repeats) and is not detected with 1 or 0 repeats [104]. This repeat region likely is susceptible to “accordion” type recombination, where there is loss as well as gain of repetitive sequences during DNA replication and recombination events. We have seen this with EBV genomes assembled in yeast and amplified in *E.coli.* It is possible this variation occurs in yeast when multiple fragments with free ends are transformed into yeast causing non-homologous end-joining and aberrant deletions. This BamW variation has also been observed during BAC recombineering in *E.coli* cells [9]. By catching these types of repeat variants early and eliminating them from our assemblies, we have, in most cases, been able to derive EBV^YA^ genomes that produce virus that can then transform primary B cells. These variations in the repeat region are also seen in the sequences of clinical isolates [105, 106] and in EBV BAC cloned genomes [9].

Another issue we observed, also reported by the Kanda lab with their Akata BAC, is the instability in the family of repeats region (FR) of oriP [55]. This was noticed in *E.coli* propagated DNA clones. Interestingly, this instability was not detected in a BAC clone of B95-8. This was attributed to the secondary structure differences between these two FR regions. We observed in our sequencing of TAR 1 that our clones had a 106 bp additional FR repeat sequence compared to the reference Akata sequence (Fig. S1a) [40]. This additional sequence in the FR region could be due to a sequencing issue or a real deletion in the particular Akata that was sequenced by the Flemington lab [40]. We designed PCR primers spanning the FR repeats to detect deletions within this region. A primer pair capable of detecting FR deletions was identified (Table S2, Fig. S2c-d). Using this assay, we screened subassemblies (TAR 1-6) and complete genomes (TAR 1-11). The majority (90%) of the assembled genomes did not contain any FR deletion (Fig. S2c); clones with detected deletions (Fig. S2c, lanes 4 and 5) were excluded from further analysis.

Our design for the EBV Akata-BX1 clone was to engineer the vector at the ends of the genome. The reason to do this was so that the vector could be excised from the EBV genome using unique restriction enzymes, in this case I-SceI. In the HSV-1 design we used Cre-Lox recombination to excise the vector. This was the plan for EBV, unfortunately we discovered a cryptic I-SceI site in TAR 11 (coordinate 166192 of Akata) and have seen the cleavage products even after limiting the digestion time. Thus, we had to be careful especially when we release the TAR 11 fragment from the vector for assembly of TAR 7-11 and subsequently for TAR 1-11. We did attempt to engineer mutations to inactivate the cryptic site, but while it blocked cleavage of a PCR product with the mutation, it did not block cleavage of the mutation in the TAR 11 backbone. We have not observed any detrimental effects on virus production and B cell transformation with the vector in the virus genome. The one potential problem could be the expression of LMP2a whose transcript would now be longer because of the 9.5 kb vector sequence. The Johannsen laboratory cloned the DFLR [59] M81 genome into a BAC vector which resides within the TR region. This vector was eliminated after extended growth passage of DFLR LCLs [59]. Future modification of the Akata-BX1^YA^ clone maybe to clone the vector in the TR region, use another restriction enzyme that is unique to this clone or excise the vector using Cre-lox.

The ability to generate HEK-293 virus producer cell lines was technically a difficult undertaking at first. We have generated many cell lines that are G418 resistant and GFP+. Not all these cell lines produce virus upon lytic induction. Our first lesson was that it was better to clone out the HEK-293 cells first rather than use pooled populations. We also screened cell lines for an increase in GFP signal upon lytic activation. The increase in GFP signal was likely due to amplification of the genome and thus an increase in copy number of CMV-GFP DNA that is transcribed. Again, not all the cells that displayed this phenotype produce infectious virus as judged by the Raji titer. Virus yields were also variable, and we had to identify and select clones that were high virus producers (Fig. 6).

One novel approach we used for virus production was to use baculovirus transduction of HEK-293 cells. Others have used plasmid transfection of *Zta*, *Rta* and *BALF4* expressing DNAs to turn on lytic gene expression [59, 78, 79]. We used an approach developed by Jeff Vieira for induction of KSHV lytic gene expression [81]. This approach used baculovirus transduction to express the KSHV replication and transcriptional activator (*RTA*) to produce virus. We similarly used BacMam vectors to express Zta, Rta and gB. Using these viruses, we could infect greater than 90% of the cells, visualized using a virus expressing Zta-mCherry (Fig. S8). Furthermore, as shown with the ΔBFRF3 mutant, we can complement this essential gene using baculovirus transduction. Hence, we can use this approach to facilitate reverse-genetic analyses of essential genes. It is possible that baculovirus transduction can also lead to induction of antiviral responses from the host cells which could mitigate transgene expression. By using the strong CMV promoter, adjusting viral titers and removal of virus after infection we can mitigate some of these antiviral impacts.

Historically, studies in our lab have investigated the capsid assembly pathways of herpesviruses. Of interest was the small capsid protein that decorates the capsid [107-110]. This protein was dispensable for HSV-1 capsid assembly in infected cells but a virus encoding a null mutation in the gene for VP26 displayed a reduction in the levels of virus production in the mouse ganglia [82]. For the betaherpesviruses, the SCP of HCMV plays an essential role that impacts major capsid protein levels in the mutant infected cells [111]. For both EBV and KSHV, we used a recombinant baculovirus expression system in insect cells to investigate the pathways of capsid assembly. Using this method, we could demonstrate the assembly of icosahedral capsids when all six capsid proteins were co-expressed in insect cells. To our surprise, we discovered that the KSHV SCP (*ORF65*) and the EBV SCP (*BFRF3*) were required for capsid assembly in insect cells [73, 83]. This was later confirmed in KSHV infected cells [112, 113]. In the present study, we find that this is also the case for EBV SCP in EBV-infected cells.

This study establishes a versatile yeast-based synthetic genomics platform for EBV by modularly assembling an infectious Akata-BX1 genome from 11 TAR-cloned fragments, demonstrating stable propagation in *E. coli*, efficient virus production from HEK-293 producer lines via BacMam-mediated lytic induction, and robust transformation of primary B cells into lymphoblastoid cell lines with careful exclusion of repeat-instability variants. Beyond recapitulating wild-type biology, the system enables rapid, localized engineering of essential functions—as illustrated by *Zta* and *BFRF3* deletion mutants whose lethal phenotypes were cleanly complemented *in trans*—highlighting the power of a “genome assembly line” that isolates problematic regions, minimizes unwanted recombination, and circumvents key limitations of traditional BAC-based EBV genetics.

## Abbreviations

EBV: Epstein-Barr virus
HSV-1: herpes simplex virus type-1
HCMV: human cytomegalovirus
KSHV: Kaposi’s sarcoma-associated herpesvirus
BL: Burkitt’s lymphoma
LCL: lymphoblastoid cell line
HEK-293: human embryonic kidney
TK: thymidine kinase
BAC: bacterial artificial chromosome
YCp: yeast centromeric plasmid
TAR: transformation-associated recombination
YA: yeast assembled
ORF: open reading frame
PCR: polymerase chain reaction
NGS: next generation sequencing
ONT: Oxford Nanopore Technologies
gDNA: genomic DNA
rBac: recombinant baculovirus
IR: internal repeat
TR: terminal repeat
FR: family of repeats
GFP: green fluorescent protein
SCP: small capsid protein
G418: geneticin
YPD: yeast peptone dextrose
HIS: histidine
URA: uracil
SD: synthetic dropout
2YT: 2X yeast extract tryptone.

## Data Summary

The supporting raw sequence data generated in this study have been deposited in Zenodo: https://doi.org/10.5281/zenodo.23921455.

GenBank accession numbers for EBV^YA^ genome nucleotide sequences:

|  |  |
|---|---|
| EBV_Akata_YA_deltaBFRF3 | QB114535 |
| EBV_Akata_YA_deltaZTA | QB114536 |
| EBV_Akata_YA_WT10 | QB114537 |
| EBV_Akata_YA_WT3 | QB114538 |

All other supporting data are available within the article and its supplementary files.

## Funding information

This work was supported by Public Health Service grant R21AI109338, R01AI137365, R01AI061382, R03AI146632, R03AI190227, and U01CA284811 from the National Institutes of Health. Normal and Oncological Tissue Collection Hub biobank is supported in part by Sidney Kimmel Comprehensive Cancer Center support grant P30 CA006973-59. Sequence analysis of Nanopore long reads was supported by UNC Vironomics Core (CA016086).

## Acknowledgements

We want to thank Lindsey Hutt-Fletcher for the Akata-BX1 cell line and the EBV Akata-BAC^LHF^ plasmid. We want to thank Joe Rizkallah, Gurjot Chand, Dylan Peters, Olivia Liao and Dev Rathod for help along the way in this long journey.

## Author Contributions

AR, KS, SC, SP, PG, SG, BP, LO, AC, JP, DD and PJD carried out the experiments. AR, KS, SC, SP, PG, SG, BP, LO, AC, JP, DD, RX, GG, SV, RFA and PJD wrote the manuscript and generated the figures.

## Conflicts of Interest

The authors declare that there are no conflicts of interest

## Ethical Approval

Human specimens were collected by the Johns Hopkins Kimmel Cancer Center Specimen Accessioning Core. Appropriate informed consent was obtained from all donors before specimen collection under a research protocol (NA00028682) approved by the Johns Hopkins Institutional Review Board.

## Notes

### Competing Interest Statement

The authors have declared no competing interest.

### Summary of Updates

New authors added New Data added Text has significant revisions

