## Supplementary material for "Modular Assembly of an Infectious Clone of the Akata strain of EBV using Synthetic Genomics Methods in Yeast": Supplementary Materials Raviv et al..pdf

**Table S1.** TAR construction primer sequences and coordinates.

| Primer | Sequence |
|---|---|
| EBVcon1R 1 | ACGGGCAAGAAGAAAAGTAAGGGTGAATAGAGCAAGACGAATTCTtagggataacagggtaatcgggtaccgagctcgaattc |
| EBVcon1F 10811 J1-2 | AATTAGTTGAGAGGCTAGTGTTTTAAACATGCACTCTAGGCCAGCtagggataacagggtaatgatcctctagagtcgacctg |
| EBVcon2R 10732 J1-2 | TCAACTAATTTCTGCGCTTGGGTTTCTAATTGGGACACATGAATTtagggataacagggtaatcgggtaccgagctcgaattc |
| EBVcon2F 35639 J2-3 | CTAAGTCACAGGCTTAGCCAGGTGATTTGTGAATTTCAGTTTATTtagggataacagggtaatgatcctctagagtcgacctg |
| EBVcon3R 35560 J2-3 | TGTGACTTAGTCTTCATCCTCTTCTTCTTCTATGTAGACAGATTGtagggataacagggtaatcgggtaccgagctcgaattc |
| EBVcon3F 54238 J3-4 | GGTCATTAGTGGCTTCGGGTAGCGTCCGATCCACGTACTCGCTCAtagggataacagggtaatgatcctctagagtcgacctg |
| EBVcon4R 54159 J3-4 | ACTAATGACCCAGGCCAGGCTAACCTGCCTCCTCCTCCAACTATCtagggataacagggtaatcgggtaccgagctcgaattc |
| EBVcon4F 69225 J4-5 | CGGTGCTCGGCGTAACCAGATTTACACCTCGGGTCTCGAGCGGAGtagggataacagggtaatgatcctctagagtcgacctg |
| EBVcon5R 69146 J4-5 | CCGAGCACCGGCCAGGCTTCCAGAGCCCAAAGCCAGAAAGATGAAtagggataacagggtaatcgggtaccgagctcgaattc |
| EBVcon5F 83589 J5-6 | TAGGTCACTTTCGGCAGAAAGATATACTTTGTTCTTTGATTTAGTtagggataacagggtaatgatcctctagagtcgacctg |
| EBVcon6R 83510 J5-6 | AAGTGACCTAGCACGACGTGCCATCAGGAAGGCTGTTTCGATCTTtagggataacagggtaatcgggtaccgagctcgaattc |
| EBVcon6F 97484 J6-7 | CCAGCTGGTCATCAGCCGCTGCGCAAACGGACTCAACGTGGTCTCtagggataacagggtaatgatcctctagagtcgacctg |
| EBVcon7R 97405 J6-7 | GACCAGCTGGTTGCTCCCATTCTTAGGTGAATTTAAGGAGGCCAGtagggataacagggtaatcgggtaccgagctcgaattc |
| EBVcon7F 115683 J7-8 | GCTCTCTCATGCGTTTGGCTACAGCATCATAGCGCTTGTTTCTGGtagggataacagggtaatgatcctctagagtcgacctg |
| EBVcon8R 115604 J7-8 | ATGAGAGAGCTATGGATGTCCACATTGACAACCAAGTGCTGAGTGtagggataacagggtaatcgggtaccgagctcgaattc |
| EBVcon8F 135983 J8-9 | CCACCACCGGCCTCCACAGACCCAGCCACCATGCTATCAGGTAACtagggataacagggtaatgatcctctagagtcgacctg |
| EBVcon9R 135905 J8-9 | CCGGTGGTGGGCTCTGAAGTGCCTTGTCCGGCTTTTGTAGGTCCGtagggataacagggtaatcgggtaccgagctcgaattc |
| EBVcon9F 148367 J9-10 | TAAATGCTGCAGTAGTAGGGATCTGGACGCGCGACCTGCTACTCTtagggataacagggtaatgatcctctagagtcgacctg |
| EBVcon10R 148288 J9-10 | GCAGCATTTACAATTTATATTATGTAAATCGGCAGCGTAGGGTACtagggataacagggtaatcgggtaccgagctcgaattc |
| EBVcon10F 162636 J10-11 | GGCATAGCGGGCAAGAAGGTTGGGCGAGAAGGAGGCCGCATAGACtagggataacagggtaatgatcctctagagtcgacctg |
| EBVcon11R 162557 J10-11 | CCGCTATGCCTACTACCTGCAGTTTTGCCAGGGACAGAAGAGCTCtagggataacagggtaatcgggtaccgagctcgaattc |
| EBVcon11F 171323 | GATGGGGGCATGGGGGGGTCGGATTTCGCCCTTATTGCCCTGTTTtagggataacagggtaatgatcctctagagtcgacctg |

*I-SceI restriction enzyme site is underlined

** EBV sequence is in uppercase and the vector sequence is in lowercase

*** Akata coordinates are shown below primer name. The 79 bp overlap sequences are from the F primer coordinates of each TAR to the R primer coordinates of the neighboring TAR. The Junctions are also shown.

**Table S2.** Junction and FR detection primer sequences.

| Primer | Sequence | PCR Size |
|---|---|---|
| Vector F | ACGACGGCCAGTGAATTG | |
| J1 R | GTGCGTGTGTACTCACAAGT | |
| J1-2 Det F | CAAAATGCGAAACTACAGGC | 561 bp |
| J1-2 Det R | ACGGGGCCACTATACTTTGC | 561 bp |
| J2-3 Det F | CACTTACAACCAAGCCACTA | 524 bp |
| J2-3 Det R | GGCGGCAACGCATTACATAG | 524 bp |
| J3-4 Det F | CTCCAGAGCCGTGTGAA | 582 bp |
| J3-4 Det R | GCGGCTACCGCCAAAGAA | 582 bp |
| J4-5 Det F | ATCTTCTTCTCCCTCAGAGTC | 553 bp |
| J4-5 Det R | CAATGATGCTGGAATGGCCC | 553 bp |
| J5-6 Det F | AATTGTTAAGCAGAGGCGTT | 572 bp |
| J5-6 Det R | CTGGCAGTCTAGGAGCCTCA | 572 bp |
| J6-7 Det F | TTAAAAGCCGTGTATTCCCC | 591 bp |
| J6-7 Det R | CAAGCTTGCGCTGGAAGATG | 591 bp |
| J7-8 Det F | TCATCGTAAAATCGAAGGGC | 657 bp |
| J7-8 Det R | ACAAGTGGGAGCCTGAACAA | 657 bp |
| J8-9 Det F | CACTTTAGTGACCTGGAACC | 547 bp |
| J8-9 Det R | CTGGGGAGGTAAAGTAGCCG | 547 bp |
| J9-10 Det F | GTACAGACGATTGTTTGGCT | 511 bp |
| J9-10 Det R | TCCGAGGGCACTCTGTAAAC | 511 bp |
| J10-11 Det F | GAAAGTTGCCCAAAAAGTCC | 531 bp |
| J10-11 Det R | ACGTGTCCACGCAGATCTTT | 531 bp |
| J11 F | TCGCCCTTATTGCCCTGTTT | |
| Vector R | CCAAGCTATTTAGGTGAGAC | |
| Vector R507 | ATTCAGGTTCATCATGCCGTTTGTG | |
| FR-F | TGCTGACTGTATATGCATGAGG | |
| FR-R | GCTGAGAGCACGGTGGAATA | |

**Table S3.** Primer sequences for cloning and mutagenesis.

| Primer Name | Sequence |
|---|---|
| *BZLF1* cDNA F XhoI | GGGG*CTCGAG*ACCATGATGGACCCAAACTCGACTTCTG |
| *BZLF1* cDNA R KpnI | GG*GGTACC*TTAGAAATTTAAGAGATCCTCGTG |
| *BZLF1mCherry* R KpnI | GG*GGTACC*CTACTTGTACAGCTCGTCCATGCC |
| *BRLF1* NheI F | GGGG*GCTAGC*ACCATGAGGCCTAAAAAGGATGGCTTG |
| *BRLF1* KpnI R | GG*GGTACC*CTAAAATAAGCTGGTGTCAAAAATAG |
| *BALF4* NotI F | GGG*GCGGCCGC*ACCATGACTCGGCGTAGGGTGCTAAGC |
| *BALF4* KpnI R | GGG*GGTACC*TTAAAACTCAGTCTCTGCCTCCCC |
| *BFRF3* XhoI F | GGGG*CTCGAG*ACCATGGCACGCCGGCTGCCCAAGCCC |
| *BFRF3* KpnI R | GG*GGTACC*CTACTGTTTCTTACGTGCCCCGCG |
| *BFRF3* gRNA | AGUACGUGCAGAGGACUUUU |
| *BFRF3* WT Det F | CATCTGTCAGCAACGCCAAG |
| *BFRF3* WT Det R | GCAGAACTGGGACGTCAGAA |
| *BFRF3* Mutant Det F | GGAGGCGGATTTTCCAGACA |
| *BFRF3* Mutant Det R | ATCGCAGACCTCGAGAGACT |
| *BZLF1* gRNA | CAACUGACUAACCAAGCCGG |
| *BZLF1* WT Det F | CTGGAAGCCACCCGATTCTT |
| *BZLF1* WT Det R | AACTGACTAACCAAGCCGGG |
| *BZLF1* Mutant Det F | TCCTCCTCTTACCCTGGGTG |
| *BZLF1* Mutant Det R | CAGATGGACCTGAACCACCC |

* Restriction enzyme sites are italicized.

**Table S4.** Junction detection nested primer sequences.

| Primer | Sequence | PCR Size |
|---|---|---|
| J1-2 Nest 1 Det F | CGGCCTTGCGAACAATCATT | 263 bp |
| J1-2 Nest 1 Det R | GATAGCACTCGACGCACTGA | 263 bp |
| J2-3 Nest 1 Det F | CCAGCGCCAATCTGTCTACA | 219 bp |
| J2-3 Nest 1 Det R | CCCTTGGCTGACCTTGGTTA | 219 bp |
| J3-4 Nest 1 Det F | TTGCCTGGGGGATAGTTGGA | 127 bp |
| J3-4 Nest 1 Det R | AAAAGACGATCAGCCGAGGC | 127 bp |
| J4-5 Nest 2 Det F | AGGTGTCGGCTGTCTTCATG | 124 bp |
| J4-5 Nest 2 Det R | CAGAAAATGCTGCGTCTCCG | 124 bp |
| J5-6 Nest 2 Det F | CCGCACGCATAACCTCAAAG | 265 bp |
| J5-6 Nest 2 Det R | AGCATACAGGGTGTTTCCGG | 265 bp |
| J6-7 Nest 3 Det F | ACATGGGGCAATTGGGCATA | 233 bp |
| J6-7 Nest 3 Det R | CTGCTTCGCTTCAGGATGGA | 233 bp |
| J7-8 Nest 3 Det F | GCAAAGCTCTGCCATAACCG | 285 bp |
| J7-8 Nest 3 Det R | CTTTGTGTTCCCTTGCTGCC | 285 bp |
| J8-9 Nest 4 Det F | TTTCTGAGAAACCGCGTGGA | 273 bp |
| J8-9 Nest 4 Det R | GTCCAGGTTGGTCTGAAGCA | 273 bp |
| J9-10 Nest 4 Det F | CGCTTTGGTTACTCGGGAGA | 219 bp |
| J9-10 Nest 4 Det R | TGGCGTTTCCGAAGAGTAGC | 219 bp |
| J10-11 Nest 4 Det F | AAGAAGAGCGTGTTCAGGCA | 327 bp |
| J10-11 Nest 4 Det R | TTTCTACATGACCCACGGCC | 327 bp |

Fig. S1

(a)

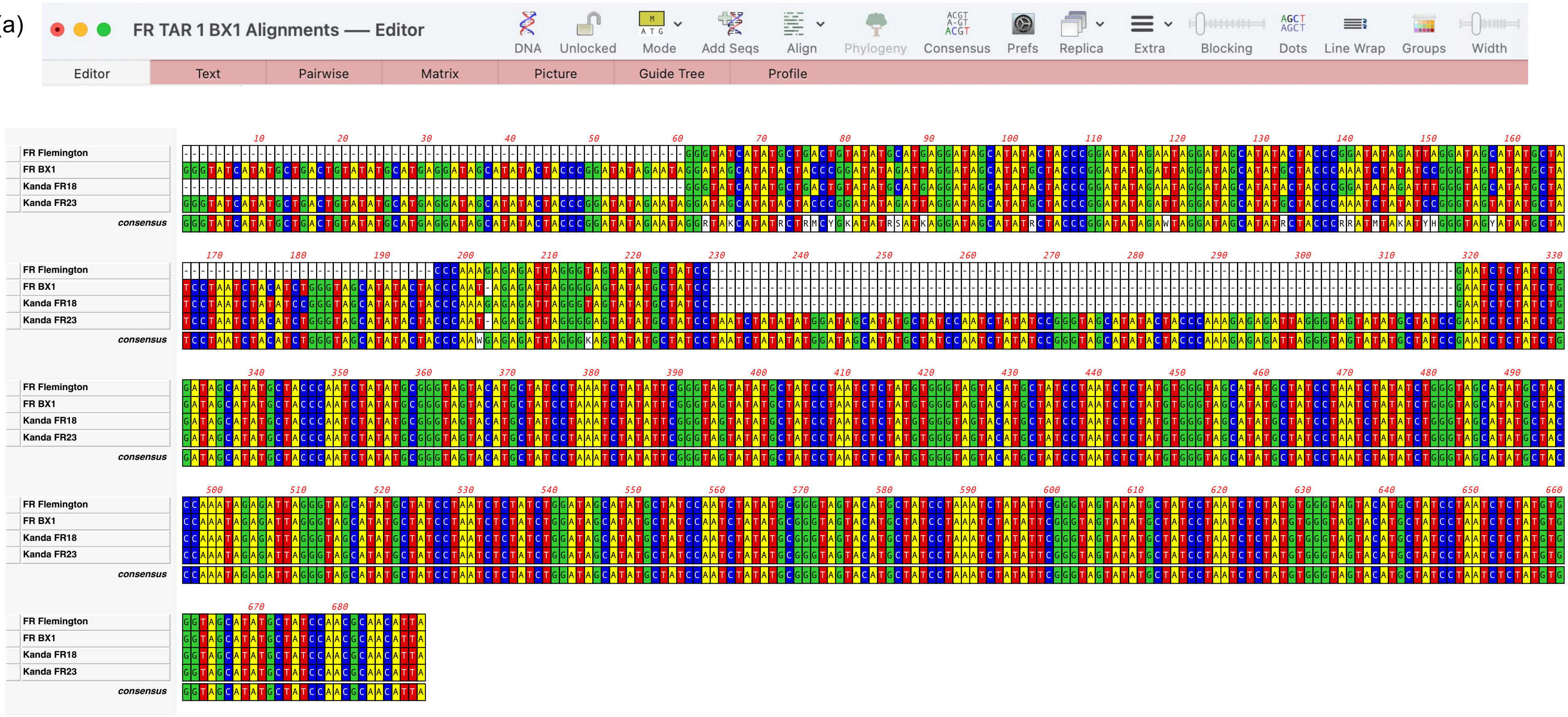
FR TAR 1 BX1 Alignments — Editor
DNA
Unlocked
Mode
Add Seqs
Align
Phylogeny
Consensus
Prefs
Replica
Extra
Blocking
Dots
Line Wrap
Groups
Width
Editor
Text
Pairwise
Matrix
Picture
Guide Tree
Profile
FR Flemington
FR BX1
Kanda FR18
Kanda FR23
consensus

Fig. S1

(b)

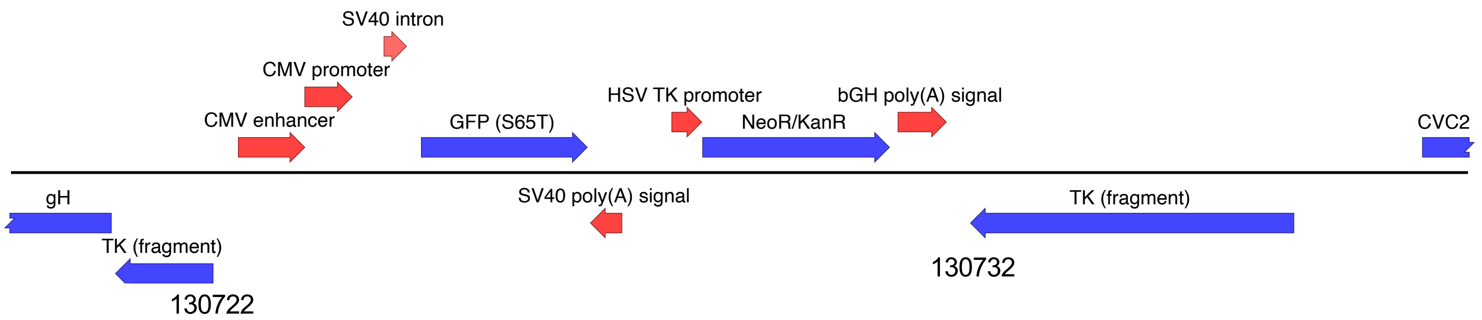

**Fig. S1.** (a) ClustlW alignment of the Family of Repeat (FR) sequences from the reference Akata genome, sequenced by the Flemington laboratory (KC207813.1)[40], from the TAR 1 plasmid cloned from Akata-BX1 and from the Akata-BAC (FR18 and FR23) generated by the Kanda laboratory [55]. (b) The genetic organization of the BX1 cassette in TAR 8. The CMV promoter drives expression of GFP and the HSV-1 TK promoter drives the neomycin gene expression. The cassette is inserted between amino acids 464 and 468 of BXLF1 (TK) which map to Akata coordinates 130732 to 130722.

Fig. S2

(a)

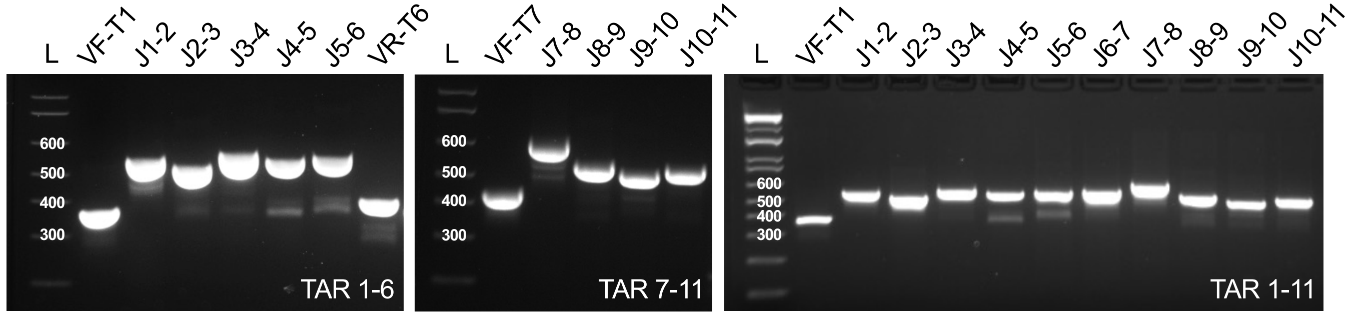

(b)

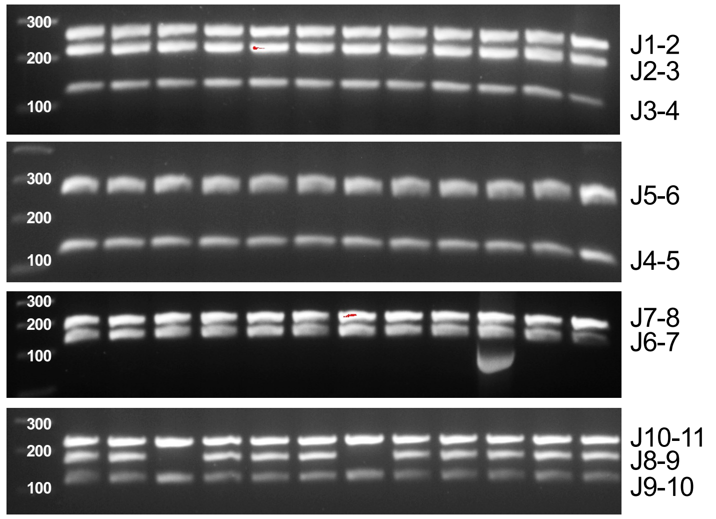

**Fig. S2.** Detection PCR analyses. (a) Detection PCR assay verifying junctions between adjacent TAR fragments following yeast assembly. Profiles are shown for sub-assemblies (TAR 1–6 and TAR 7–11) and the complete genome (TAR 1–11). Junction nomenclature corresponds to the flanking fragments (e.g., J1-2 detects the linkage between TAR 1 and TAR 2); 'V' denotes the vector junction. DNA molecular weight standards (base pairs, bp) are indicated on the left (L, DNA ladder). (b) Nested PCR detection assays used for rapid multi-junction screening of different TAR 1-11 assembled genomes. (c) PCR-based differentiation of FR repeat length in different genome assemblies. (d) PCR detection of the FR region in yeast assembled genomes (WT and ΔFR) compared to genomic Akata and Akata-BX1 DNA (isolated from both uninduced and lytic induced {Ind} cells) and Akata-BAC$^{LHF}$ DNA. DNA molecular weight standards (bp) are shown on the left (L).

(c)

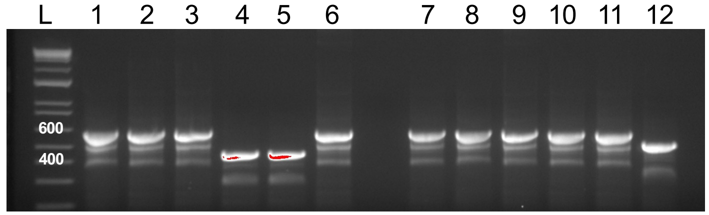

(d)

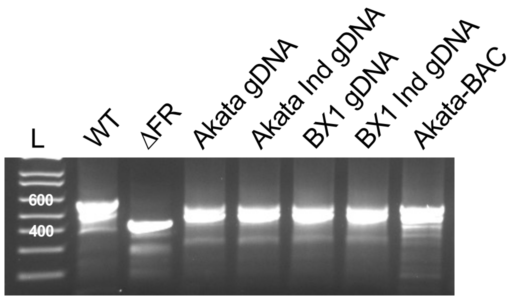

Fig. S3

(a)

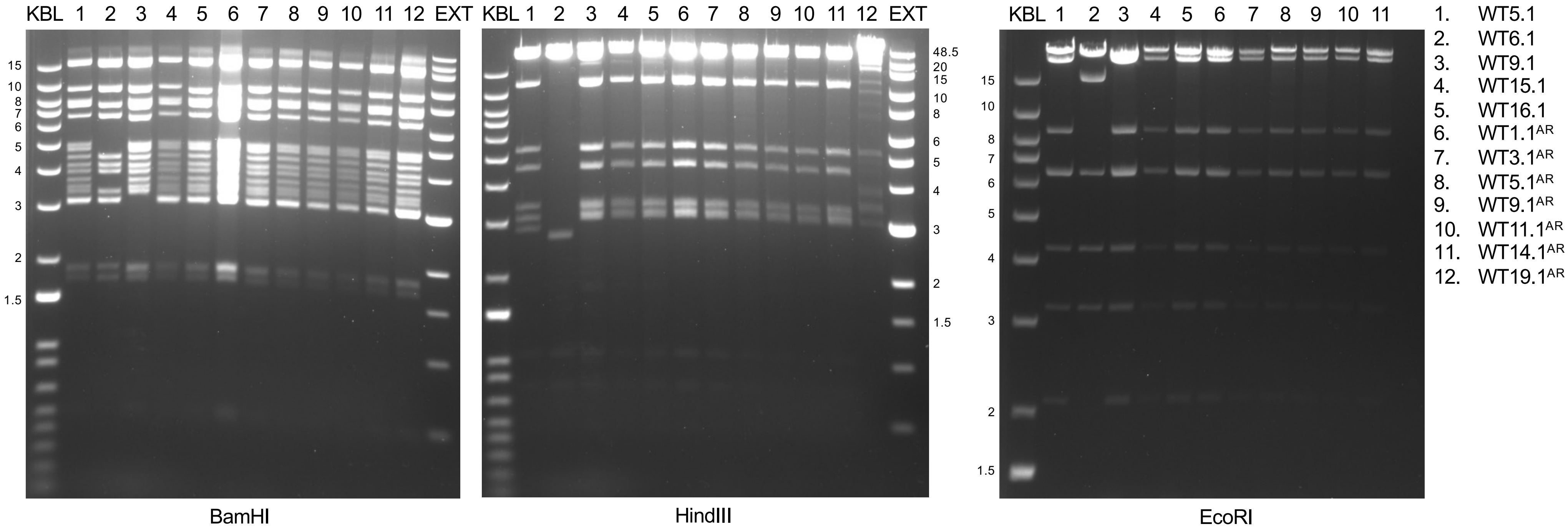
KBL
1
2
3
4
5
6
7
8
9
10
11
12
EXT
15
10
8
7
6
5
4
3
2
1.5
BamHI
48.5
20
HindIII
EcoRI
1. WT5.1
2. WT6.1
3. WT9.1
4. WT15.1
5. WT16.1
6. WT1.1AR
7. WT3.1AR
8. WT5.1AR
9. WT9.1AR
10. WT11.1AR
11. WT14.1AR
12. WT19.1AR

Fig. S3

(b)

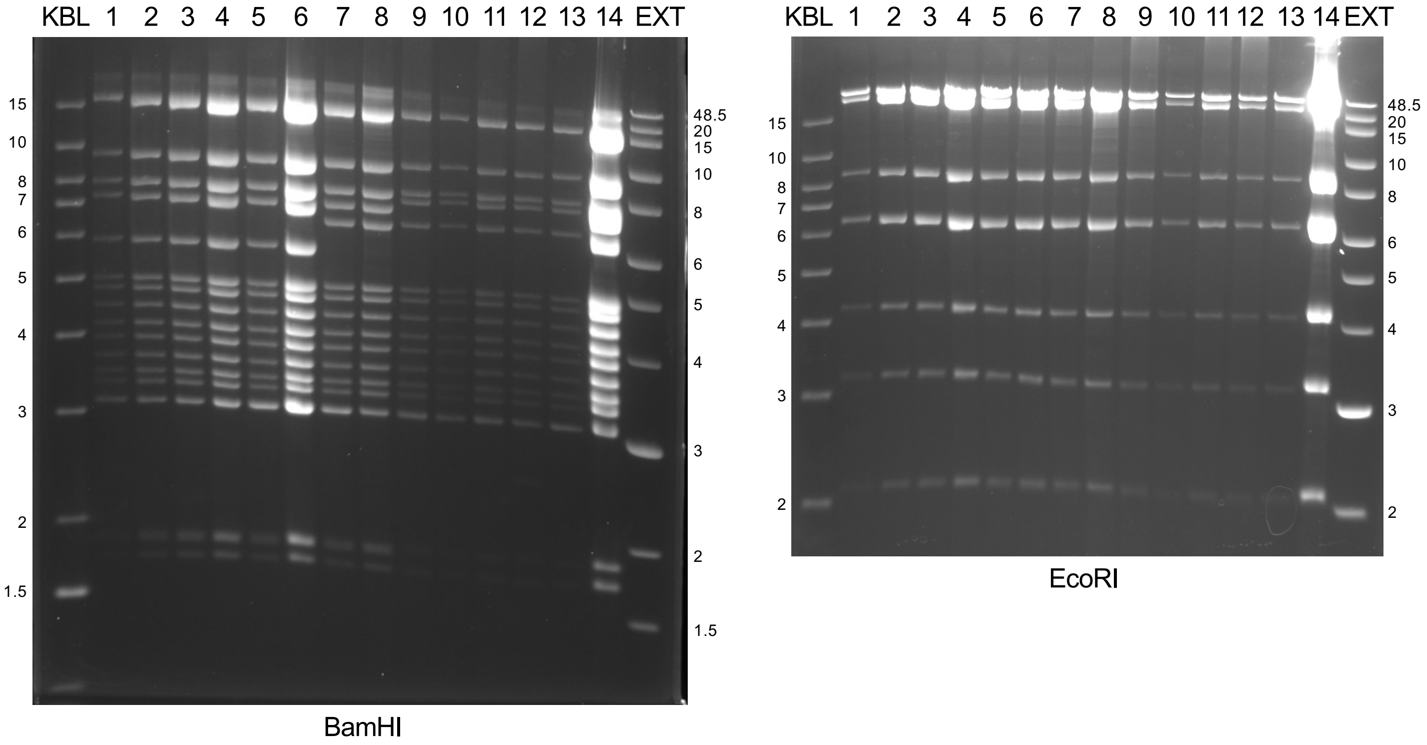

1. ΔBFRF3 11.2.1
2. ΔBFRF3 11.6.1
3. ΔBFRF3 11.9.1
4. ΔBFRF3 11.13.1
5. ΔBFRF3 11.14.1
6. ΔBFRF3 11.15.1
7. ΔBFRF3 14.10.1
8. ΔBFRF3 14.30.1
9. ΔZTA 3.1
10. ΔZTA 4.1
11. ΔZTA 7.1
12. ΔZTA 29.1
13. ΔZTA 29.2
14. ΔZTA 31.1

Fig. S3

(c)

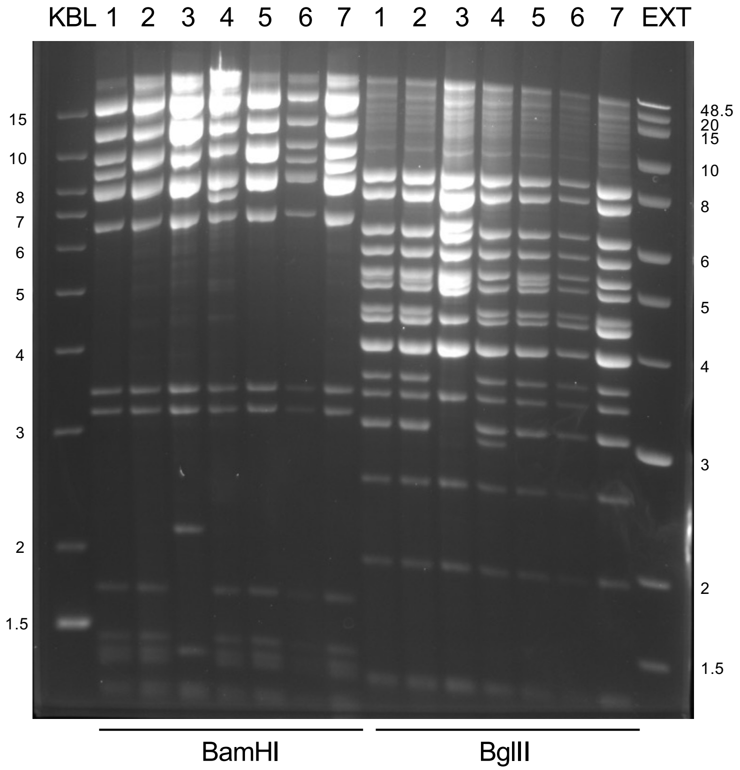

1. T7-11 5
2. T7-11 BX1 36.2
3. T7-11 hrGFP 39.2.4
4. T7-11 4.2
5. T7-11 BX1 36.2
6. T7-11 5
7. T7-11 5

**Fig. S3.** Analysis of TAR 1-6 and TAR 7-11 assemblies. (a) Independent clones of TAR 1-6 sub-genomes of wild-type (WT) were analyzed by restriction enzyme digestion (BamHI, HindIII and EcoRI). (b) Similarly, TAR 1-6 clones of ΔZta and ΔBFRF3 were analyzed by BamHI and EcoRI restriction enzyme digestion. (c) WT TAR 7-11 sub-genomes (clones 5, BX1, 4.2) and hrGFP39 were checked by digestion with BamHI and BglII. The digests were run on 0.9% TAE gels at low voltage (35-50V). The gels were imaged at different times to capture the resolution of both small and large fragments. DNA standards are shown in lane KBL (1kb + ladder) and EXT (1 kb extend ladder). The sizes (kb) of the standards are shown for reference.

### Fig. S4

(a)

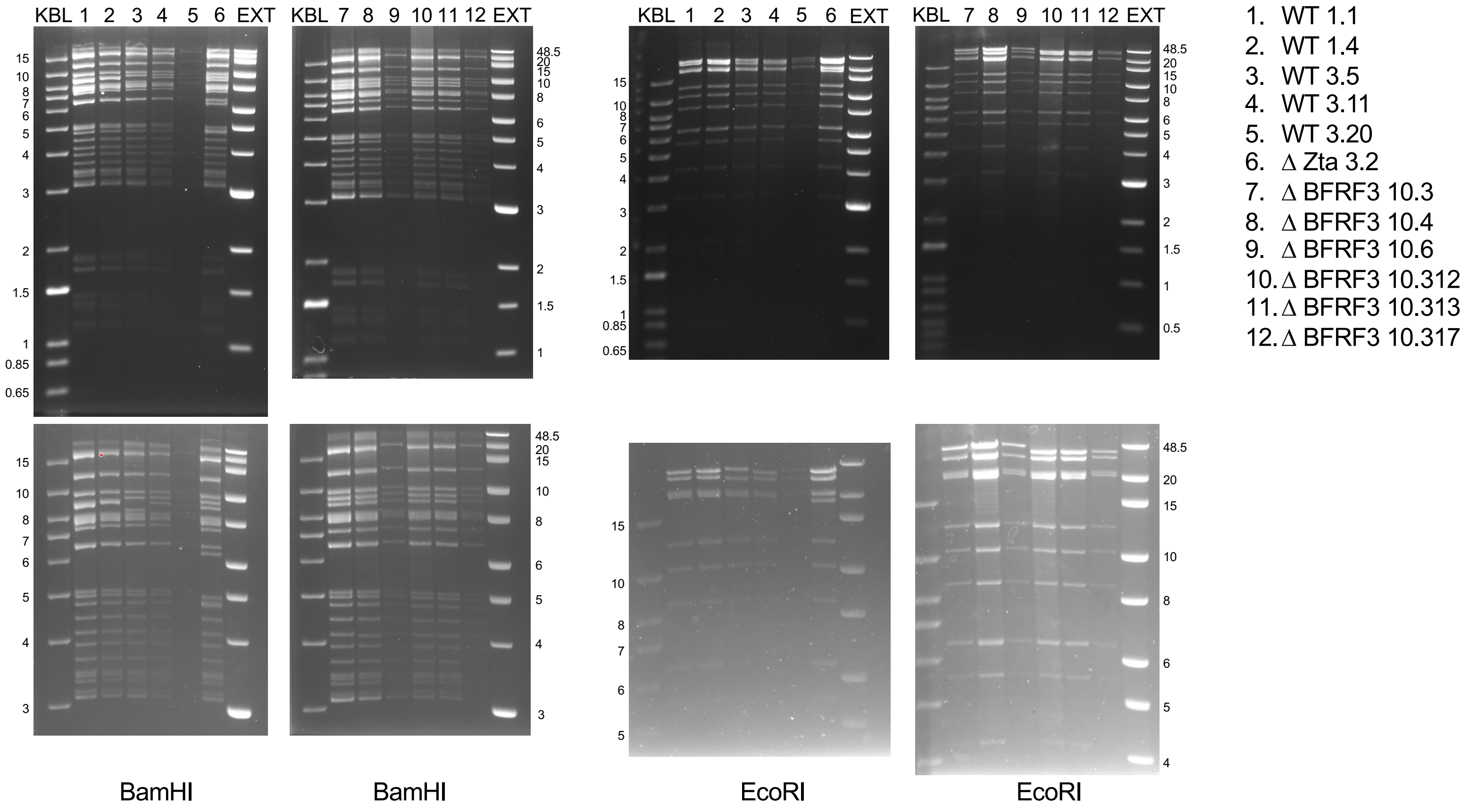

KBL 1 2 3 4 5 6 EXT
KBL 7 8 9 10 11 12 EXT
KBL 1 2 3 4 5 6 EXT
KBL 7 8 9 10 11 12 EXT
1. WT 1.1
2. WT 1.4
3. WT 3.5
4. WT 3.11
5. WT 3.20
6. Δ Zta 3.2
7. Δ BFRF3 10.3
8. Δ BFRF3 10.4
9. Δ BFRF3 10.6
10. Δ BFRF3 10.312
11. Δ BFRF3 10.313
12. Δ BFRF3 10.317
BamHI
BamHI
EcoRI
EcoRI

Fig. S4

(b)

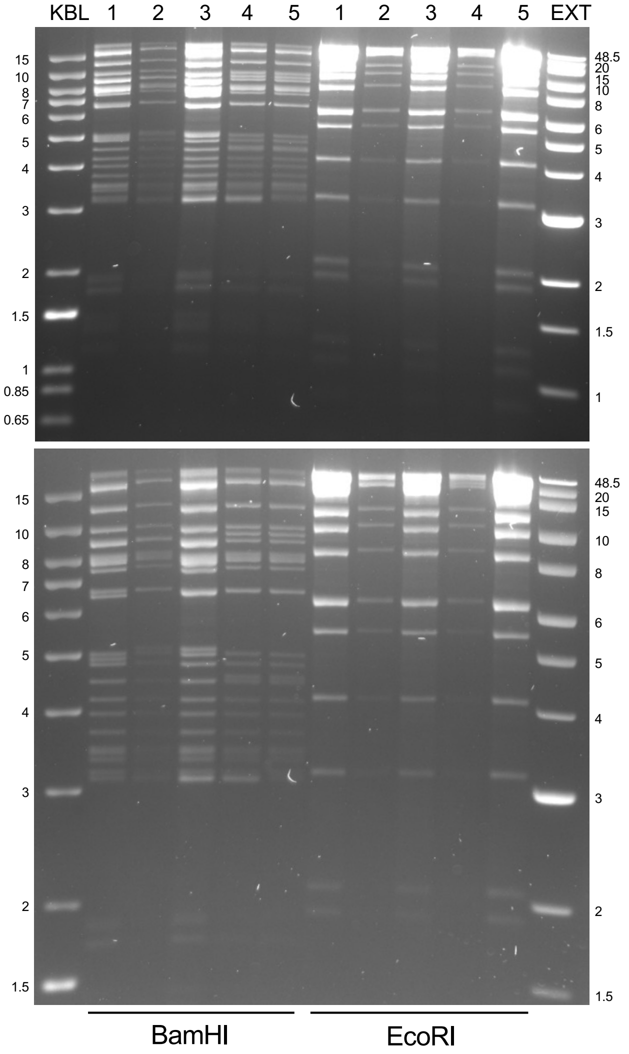

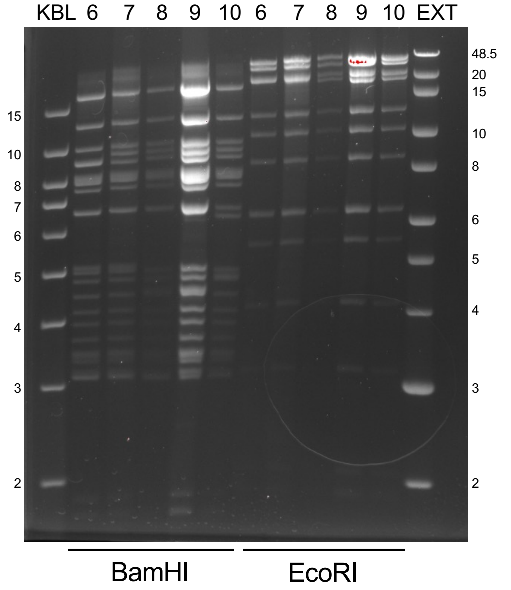

1. WT 16.7
2. WT 16.12
3. WT 16.20
4. ∆ Zta 4.7
5. ∆ Zta 7.4
6. WT 16.21
7. WT 10.1
8. ∆ Zta 4.7
9. ∆ Zta 7.4
10. ∆ Zta 3.2

**Fig. S4.** Analysis of TAR 1-11 genome assemblies. (a) Independent clones of full-length genomes of wild-type (WT), ∆Zta and ∆BFRF3 were analyzed by BamHI and EcoRI restriction enzyme digestion. (b) Similarly, DNA of clones of WT and ∆Zta were digested with BamHI and EcoRI. The digests were run on 0.9% TAE gels at low voltage (35-50V). The gels were imaged at different times to capture the resolution of both small and large fragments. DNA ladders are shown in lane KBL (1kb + ladder) and EXT (1 kb extend ladder). The sizes (kb) of the standards are indicated.

Fig. S5

(a)

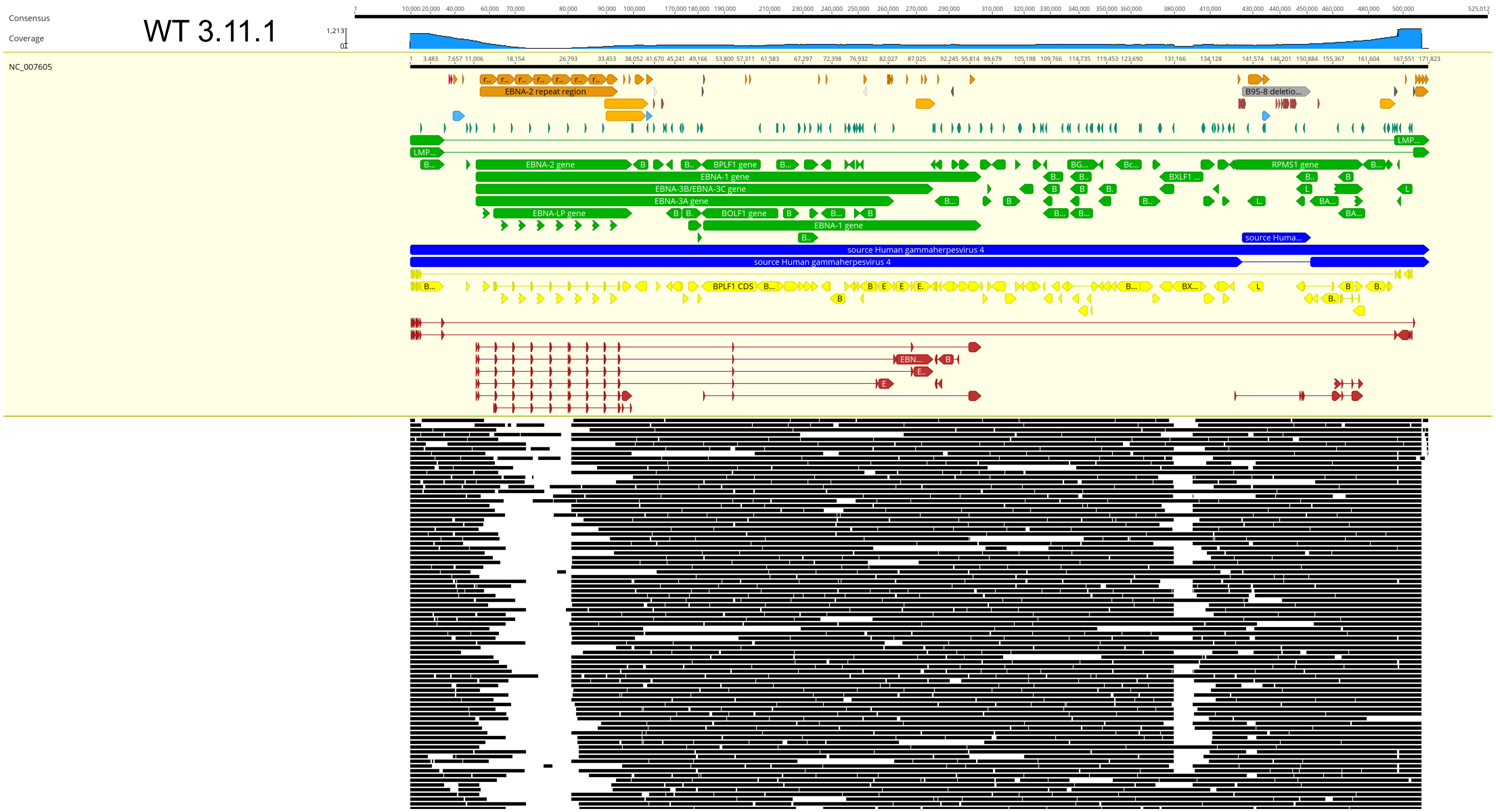

WT 3.11.1
Consensus
Coverage
NC_007605
EBNA-2 repeat region
B95-8 deletio...
EBNA-2 gene
BPLF1 gene
RPMS1 gene
EBNA-1 gene
EBNA-3B/EBNA-3C gene
EBNA-3A gene
EBNA-LP gene
BOLF1 gene
source Human gammaherpesvirus 4
BPLF1 CDS

Fig. S5

(b)

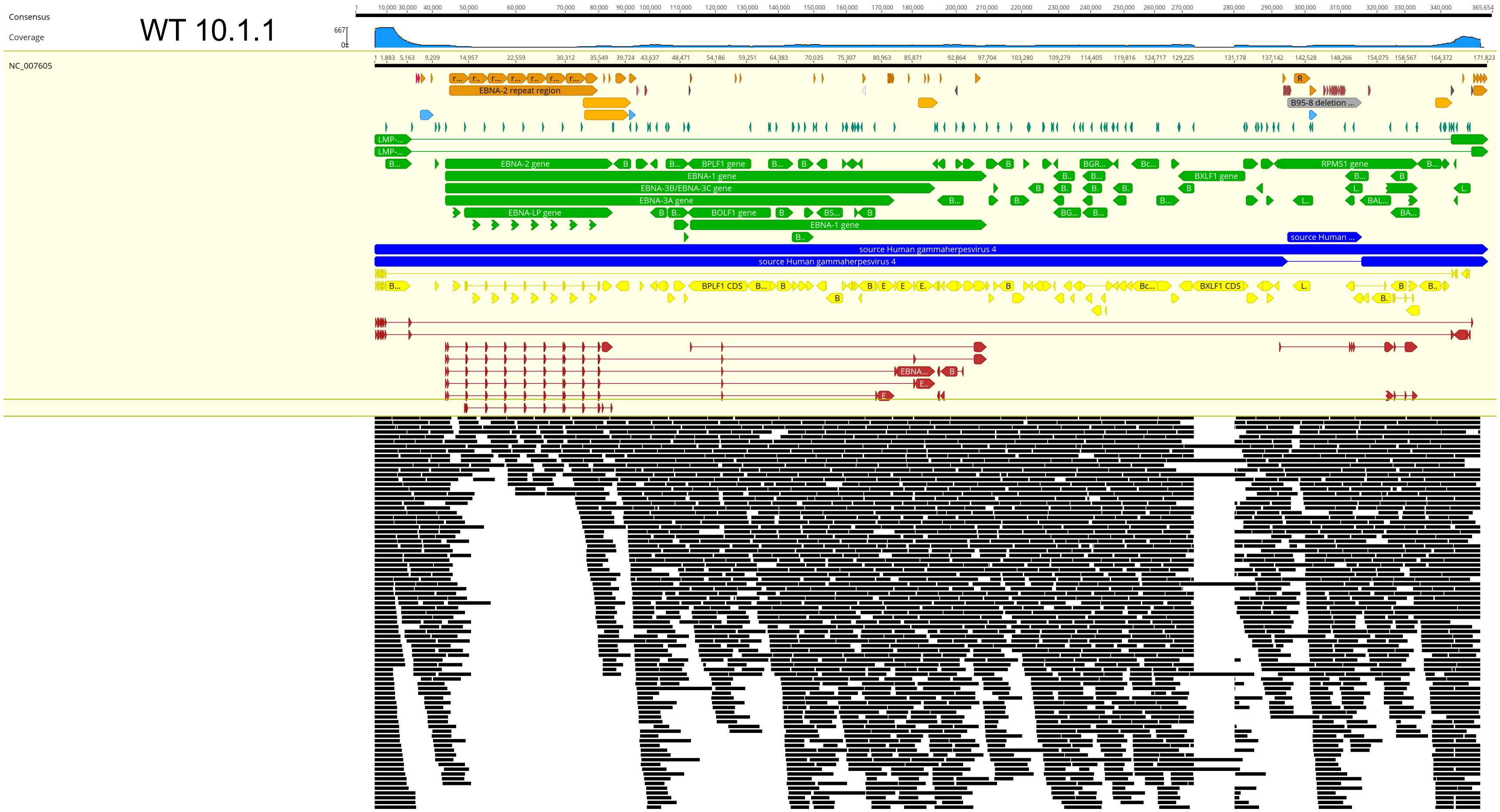

**Fig. S5**. Oxford Nanopore sequencing of EBV$^{YA}$ genomes. Sequencing was carried out by the MGH CCIB DNA Core. Long-reads for (a) WT 3.11.1 and (b) WT 10.1.1 were generated and aligned to the EBV reference genome (NC_007605) using minimap2 as implemented in Genious (Dotmatics, Inc.). Each black bar represents a single read.

### Fig. S6

(a)

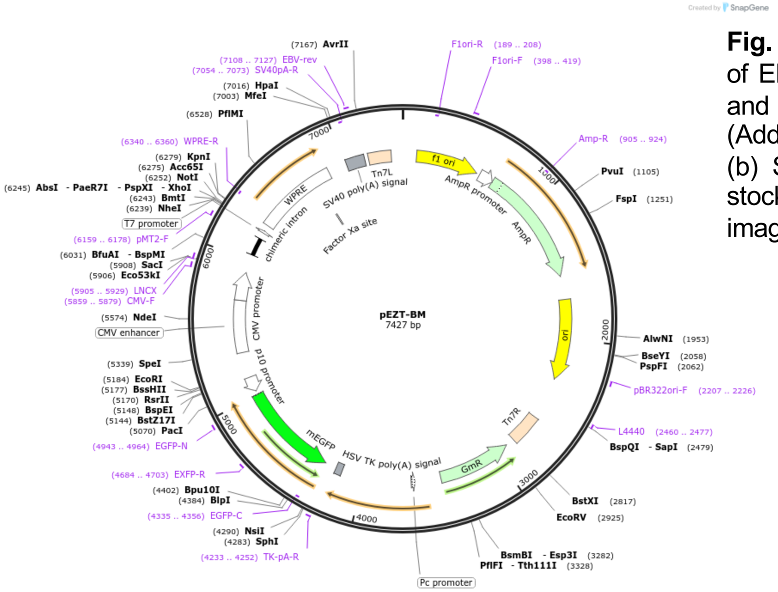

**Fig. S6.** BacMam baculoviruses for lytic induction of EBV. (a) The open reading frames for Zta, Rta and BALF4 (gB) were cloned into pEZT-BM (Addgene plasmid #74099 Ryan Hibbs laboratory). (b) Sf21 insect cells were infected with working stocks of the baculoviruses and the fluorescence imaged 48 h post-infection. Scale bar = 100 μm.

(b)

Sf21-rBac Zta Sf21-rBac Rta Sf21-rBac gB

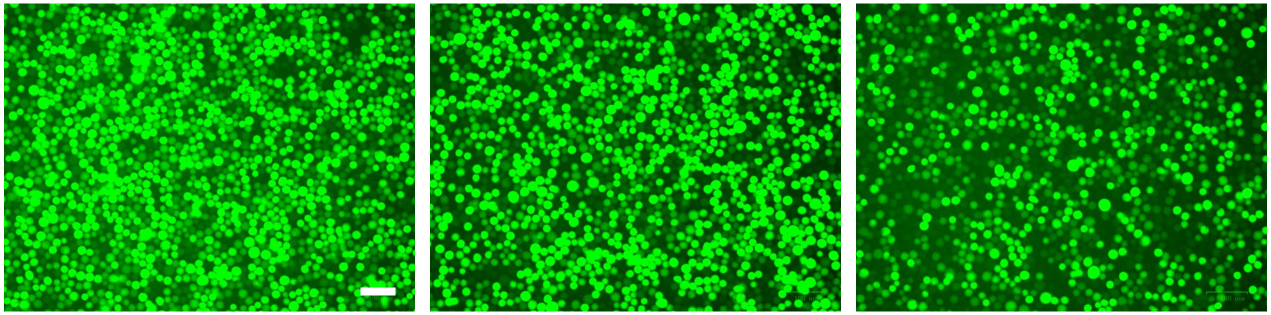

### Fig. S7

(a)

#### pEZT BZLF1

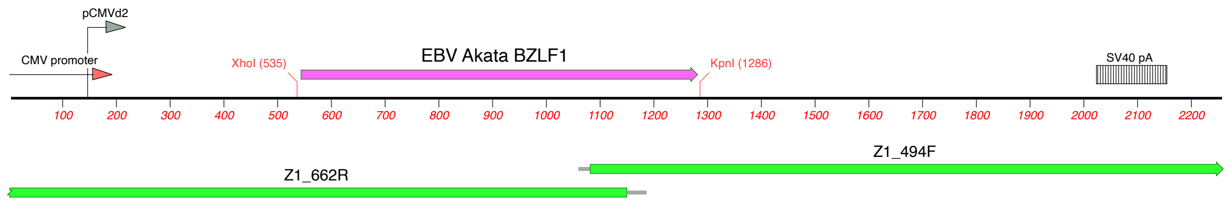

ATGATGGACCCAAACTCGACTTCTGAAGATGTAAAATTTACACCTGACCCATACCAGGTGCCTTTTGTACAAGCTTTTGACCAAGCTACCAGAGT
CTATCAGGACCTGGGAGGGCCATCACAAGCTCCTTTGCCTTGTGTGMTGTGGCCGGTGCTGCCAGAGCCTCTGCCACAAGGCCAGCTCACTGCCT
ATCATGTTTCAGCCGCACCAACTGGGTCGTGGTTTCCTGCCCCTCAGCCTGCTCCTGAGAATGCTTATCAAGCTTATGCAGCACCTCAGCTGTTC
CCAGTCTCAGACATAACCCAGAATCAACTGACTAACCAAGCCGGGGGAGAAGCACCTCAACCTGGAGACAATTCTACTGTTCAACCAGCAGCAGC
AGTGGTGTTGGCTTGCCCCGGGGCTAACCAAGAACAACAGCTAGCAGACATTGGTGCTCCACAGCCTGCACCAGCGGCTGCCCCGGCACGACGTA
CAAGGAAACCACTACAGCCAGAATCGCTGGAGGAATGCGATTCTGAACTAGAAATAAAGCGATACAAGAATCGGGTGGCTTCCAGAAAATGCCGG
GCCAAGTTTAAGCATCTGCTGCAGCACTACCGTGAGGTGGCTTCTGCCAAATCATCTGAAAATGACAGGCTGCGCCTCCTGTTGAAGCAGATGTG
CCCAAGCCTGGATGTTGACTCCATTATCCCCCGGACACCAGATGTTTTACACGAGGATCTCTTAAATTTCTAA

#### Fig. S7

(b)

pEZT BRLF1

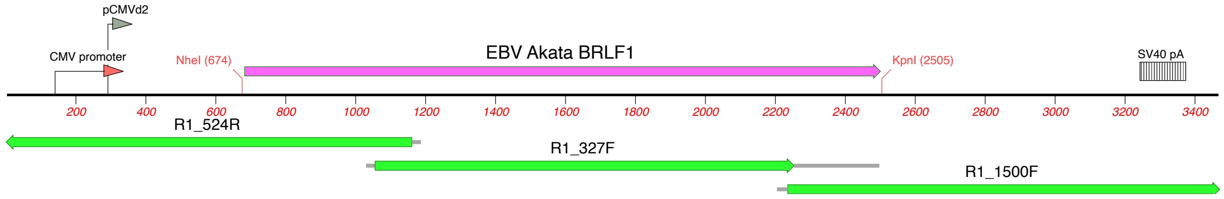

ATGAGGCCTAAAAAGGATGGCTTGGAAGACTTTCTGAGGCTAACTCCTGAAATCAAAAAGCAGCTGGGCTCTCTGGTCTCTGACTACTGCAACG
TCCTCAACAAGGAATTTACAGCCGGGAGTGTGGAGATTACTCTGAGATCCTACAAAATATGCAAGGCATTTATAAATGAGGCCAAGGCCCACGG
GCGAGAATGGGGCGGGCTAATGGCCACGCTCAACATCTGCAATTTTTGGGCCATTCTCCGAAACAACAGGGTAAGAAGACGGGCTGAGAATGCC
GGCAACGACGCATGTTCCATCGCGTGCCCTATAGTGATGCGCTACGTGTTAGACCACCTGATAGTGGTCACTGACAGATTCTTCATCCAGGCCC
CCAGCAACCGGGTGATGATTCCTGCCACCATAGGCACCGCTATGTACAAGCTCCTAAAACACAGTCGGGTGCGGGCCTACACCTACAGCAAGGT
GCTGGGCGTGGACCGCGCGGCCATCATGGCCTCCGGCAAGCAGGTAGTGGAACACCTGAACAGGATGGAGAAGGAAGGCCTCCTAAGCTCCAAG
TTCAAGGCCTTTTGCAAGTGGGTGTTCACCTATCCCGTCCTCGAGGAGATGTTCCAGACTATGGTCTCGTCCAAGACAGGCCATCTGACGGACG
ATGTTAAGGATGTCAGGGCTCTGATTAAGACACTGCCCCGGGCCTCCTACTCCAGCCACGCCGGACAGAGGAGCTACGTGAGCGGCGTGCTTCC
CGCGTGCCTGCTGTCAACCAAGTCCAAGGCAGTGGAAACTCCTATCCTCGTGTCTGGAGCCGACAGGATGGACGAGGAGCTCATGGGGAATGAT
GGGGGTGCCTCTCACACCGAGGCCCGCTACTCGGAGTCCGGACAGTTTCATGCTTTTACAGATGAACTCGAAAGTCTCCCGAGCCCGACCATGC
CCCTGAAGCCCGGTGCCCAAAGCGCCGACTGCGGTGACAGCAGTTCCAGCAGCAGTGACTCGGGCAACAGTGACACCGAGCAGAGCGAGCGGGA
AGAGGCCAGGGCCGAGGCCCCGCGCCTGCGGGCCCCAAAGTCGCGCCGGACATCCAGGCCCAACCGTGGTCAAACTCCATGTCCTTCCAACGCG
GAGGAACCTGAACAGCCTTGGATAGCAGCGGTCCACCAAGAGAGCGATGAGAGACCCATATTCCCCCACCCCTCAAAGCCCACCTTTCTTCCTC
CCGTTAAAAGGAAGAAGGGCCTCAGGGATAGCCGGGAAGGTATGTTCCTGCCAAAGCCGGAAGCGGGCAGTGCCATATCTGACGTGTTCGAGGG
GCGAGAGGTGTGTCAGCCAAAGAGGATCAGGCCCTTCCATCCACCCGGATCCCCGTGGGCCAACCGGCCCCTGCCTGCCTCTTTGGCTCCCACC
CCCACAGGACCTGTCCATGAACCGGTCGGATCCCTAACGCCAGCCCCGGTGCCCAAGCCACTTGACCCGGCCCCCGCAGTAACCCCCGAGGCAA
GTCATCTGTTGGAGGACCCTGATGAGGAAACCAGTCAGGCCGTGAAGGCCCTAAGGGAGATGGCTGACACTGTTATTCCCCAGAAGGAGGAAGC
AGCCATATGTGGACAGATGGACCTGAACCACCCGCCCCCTCGTGGCCATTTGGACGAACTGACCACAACACTAGAGTCCATGACAGAGGATTTG
AATCTGGACTCCCCCCTGACCCCAGAACTTAATGAAATCTTGGATACATTTCTAAATGATGAATGTCTGCTGCATGCCATGCATATTTCAACTG
GGCTGTCTATTTTTGACACCAGCTTATTTTAG

Fig. S7

(c)

pEZT BALF4

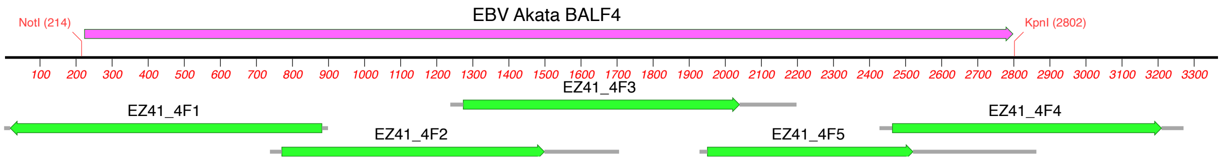

ATGACTCGGCGTAGGGTGCTAAGCGTGGTCGTGCTGCTAGCCGCCCTGGCGTGCCGCCTCGGTGCGCAGACCCCAGAGCAGCCCGCGCCCCCC
GCCACCACGGTGCAGCCTACCGCCACGCGTCAGCAAACCAGCTTTCCTTTCCGAGTCTGCGAGCTCTCCAGCCACGGCGACCTGTTCCGCTTC
TCCTCGGACATCCAGTGTCCCTCGTTTGGCACGCGGGAGAATCACACGGAGGGCCTGTTGATGGTGTTTAAAGACAACATTATTCCCTACTCG
TTTAAGGTCCGCTCCTACACCAAGATAGTGACCAACATTCTCATCTACAATGGCTGGTACGCGGACTCCGTGACCAACCGGCACGAGGAGAAG
TTCTCCGTTGACAGCTACGAAACTGACCAGATGGATACCATCTACCAGTGCTACAACGCGGTCAAGATGACAAAAGATGGGCTGACGCGCGTG
TATGTAGACCGCGACGGAGTTAACATCACCGTCAACCTAAAGCCCACCGGGGGCCTGGCCAACGGGGTGCGCCGCTACGCCAGCCAGACGGAG
CTCTATGACGCCCCCGGGTGGTTGATATGGACTTACAGAACAAGAACTACCGTCAACTGCCTGATAACTGACATGATGGCCAAGTCCAACAGC
CCCTTCGACTTCTTTGTGACCACCACCGGGCAGACTGTGGAAATGTCCCCTTTCTATGACGGGAAAAATAAGGAAACCTTCCATGAGCGGGCA
GACTCCTTCCACGTGAGAACTAACTACAAGATAGTGGACTACGACAACCGAGGGACGAACCCGCAAGGCGAACGCCGAGCCTTCCTGGACAAG
GGCACTTACACCCTATCTTGGAAGCTCGAGAACAGGACAGCCTACTGCCCGCTTCAACACTGGCAAACCTTTGACTCGACCATCGCCACAGAA
ACAGGGAAGTCAATACATTTTGTGACTGACGAGGGCACCTCTAGCTTCGTGACCAACACAACCGTGGGCATAGAGCTCCCGGACGCCTTCAAG
TGCATCGAAGAGCAGGTGAACAAGACCATGCATGAAAAGTACGAGGCCGTCCAGGATCGTTACACGAAGGGCCAGGAAGCCATTACATATTTT
ATAACGAGCGGAGGATTGTTATTAGCTTGGCTACCTCTGACCCCGCGCTCGTTGGCCACCGTCAAGAACCTGACGGAGCTTACCACTCCGACT
TCCTCACCCCCCAGCAGTCCATCGCCCCCCGCCCCACCCGCGGCCCGCGGGAGCACCTCCGCCGCCGTTCTGAGGCGCCGGAGGCGGGATGCG
GGGAATGCCACCACACCGGTGCCCCCCGCGGCCCCCGGGAAGTCCCTGGGCACCCTCAACAATCCCGCCACCGTCCAGATCCAATTTGCCTAC
GATTCCCTGCGCCGCCAGATCAACCGCATGCTGGGAGACCTCGCGCGGGCCTGGTGCCTGGAGCAGAAGAGGCAGAACATGGTGCTGAGAGAA
CTAACCAAGATTAATCCCACCACCGTCATGTCCAGCATCTACGGTAAGGCGGTGGCGGCCAAGCGCCTGGGGGATGTCATCTCAGTCTCCCAG
TGCGTGCCCGTTAACCAGGCCACCGTCACCCTGCGCAAGAGCATGAGGGTCCCTGGCTCCGAGACCATGTGCTACTCGCGCCCCCTGGTGTCC
TTCAGCTTTATCAACGACACCAAGACCTACGAGGGACAGCTGGGCACCGACAACGAGATCTTCCTCACAAAAAAGATGACGGAGGTGTGCCAG
GCGACCAGCCAGTACTACTTCCAGTCCGGCAACGAGATCCACGTTTACAACGACTACCACCACTTTAAAACCATCGAGCTGGACGGCATTGCC
ACCCTGCAGACCTTCATCTCACTAAACACCTCCCTCATCGAGAACATTGACTTTGCCTCCCTGGAGCTGTACTCACGGGACGAACAGCGTGCC
TCCAACGTCTTTGACCTGGAGGGCATCTTCCGGGAGTACAACTTCCAGGCGCAAAACATCGCCGGCCTGCGGAAGGATTTGGACAATGCAGTA
TCAAACGGAAGAAATCAATTCGTGGACGGCCTGGGGGAACTTATGGACAGTCTGGGTAGCGTGGGTCAGTCCATCACCAACCTAGTCAGCACG
GTGGGGGGTTTGTTTAGCAGCCTGGTCTCTGGTTTCATCTCCTTCTTCAAAAACCCCTTCGGCGGCATGCTCATTCTGGTCCTGGTGGCGGGG
GTGGTGATCCTGGTTATTTCCCTCACGAGGCGCACGCGCCAGATGTCGCAGCAGCCGGTGCAGATGCTCTACCCCGGGATCGACGAGCTCGCT
CAGCAACATGCCTCTGGTGAGGGTCCAGGCATTAATCCCATTAGTAAGACAGAATTACAAGCCATCATGTTAGCGCTGCATGAGCAAAACCAG
GAGCAAAAGAGAGCAGCTCAGAGGGCGGCCGGACCCTCAGTGGCCAGCAGAGCATTGCAGGCAGCCAGGGACCGTTTTCCAGGCCTACGCAGA
AGACGCTATCACGATCCAGAGACCGCCGCCGCACTGCTTGGGGAGGCAGAGACTGAGTTTTAA

### Fig. S7

(d)

pEZT BFRF3

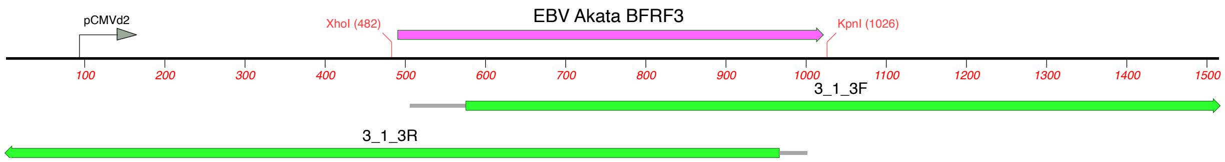

ATGGCACGCCGGCTGCCCAAGCCCACCCTCCAGGGGAGGCTGGAGGCGGATTTTCCAGACAGTCCCCTGCTTCCTAAATTTCAAGAGCTGAACC
AGAATAATCTCCCCAATGATGTTTTTCGGGAGGCTCAAAGGAGTTACCTGGTATTTCTGACGTCCCAGTTCTGCTACGAAGAGTACGTGCAGAG
GACTTTTGGGGTGCCTCGGCGCCAACGCGCCATAGACAAGAGGCAGAGAGCCAGTGTGGCTGGGGCTGGTGCTCATGCACACCTTGGCGGGTCA
TCCGCCACCCCCGTCCAGCAGGCTCAGGCCGCCGCATCCGCTGGGACTGGGGCCTTGGCATCATCAGCGCCGTCCACGGCCGTAGCCCAGTCCG
CGACCCCCTCTGTTTCTTCATCTATTAGCAGCCTCCGGGCCGCGACTTCGGGGGCGACTGCCGCCGCCTCTGCCGCCGCAGCCGTCGATACCGG
GTCAGGTGGCGGGGGACAACCCCAAGACACCGCCCCGCGCGGGGCACGTAAGAAACAGTAG

**Fig. S7.** BacMam cloning of EBV genes for lytic induction and complementation. The (a) BZLF1 (Zta), (b) BRLF1 (Rta), (c) BALF4 (gB) and (d) BFRF3 open reading frames (pink) were cloned into the baculovirus transfer vector, pEZT. The genes were PCR amplified and then cloned using the restriction enzyme sites shown in red. The cloned genes were sequence confirmed using Sanger sequencing (green) and the consensus sequence derived of the gene is shown below the cartoon. The CMV promoter elements and the SV40 polyA sequence are shown for some sequence annotations.

Fig. S8

HEK-293 rBac-Zta-mCherry

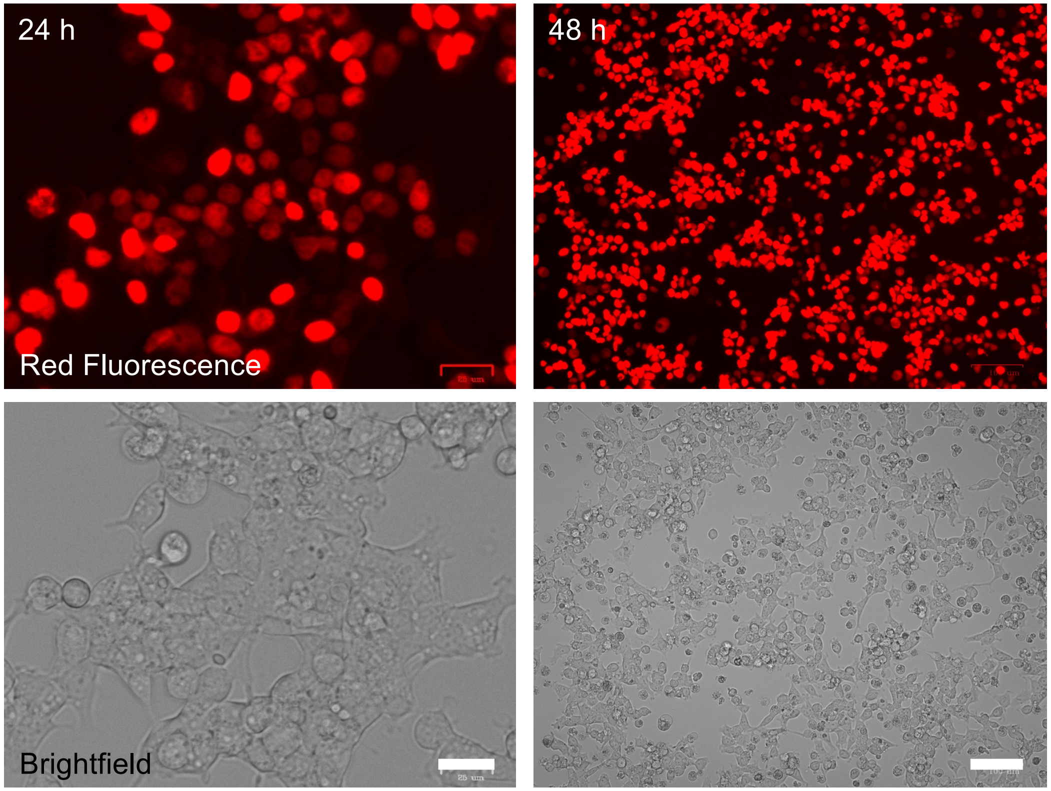

**Fig. S8.** Baculovirus transduction of human epithelial cells. HEK-293 monolayers were infected with a baculovirus expressing a Zta-mCherry fusion protein. The infected cells were imaged at 24 and 48 hours post-infection. Scale bar = 25 μm (24 h) and 100 μm (48 h).

Fig. S9

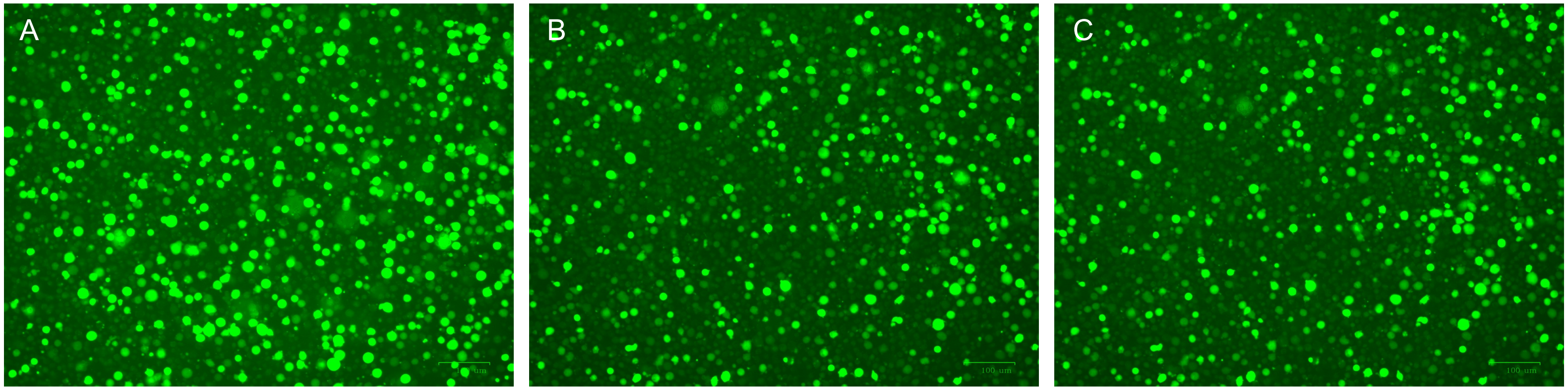

2.13 x $10^5$ RGU/ml 1.23 x $10^5$ RGU/ml 9.9 x $10^4$ RGU/ml

**Fig. S9**. Raji titration of $EBV^{YA}$ virus. Three (A, B and C) different virus preparations were made from $EBV^{YA}$ WT HEK-293 cell line. The virus in the supernatant culture medium (4.5 ml) was concentrated by centrifugation and the pelleted virus resuspended in 100 µl of PBS. The total volume of the concentrated virus plus residual media was 150 µl. 2 x $10^6$ million Raji cells were infected with 50 µl of this virus stock and imaged for GFP fluorescence 48 h post-infection. The Raji cells were serially diluted and the GFP positive cells counted (three replicate counts). The titers (Raji GFP Units, RGU) determined are shown for each virus preparation. Scale bar = 100 µm.
